# Statistical inference of the rates of cell proliferation and phenotypic switching in cancer

**DOI:** 10.1101/2022.08.31.505619

**Authors:** Einar Bjarki Gunnarsson, Jasmine Foo, Kevin Leder

## Abstract

Recent evidence suggests that nongenetic (epigenetic) mechanisms play an important role at all stages of cancer evolution. In many cancers, these mechanisms have been observed to induce dynamic switching between two or more cell states, which commonly show differential responses to drug treatments. To understand how these cancers evolve over time, and how they respond to treatment, we need to understand the state-dependent rates of cell proliferation and phenotypic switching. In this work, we propose a rigorous statistical framework for estimating these parameters, using data from commonly performed cell line experiments, where phenotypes are sorted and expanded in culture. The framework explicitly models the stochastic dynamics of cell division, cell death and phenotypic switching, and it provides likelihood-based confidence intervals for the model parameters. The input data can be either the fraction of cells or the number of cells in each state at one or more time points. Through a combination of theoretical analysis and numerical simulations, we show that when cell fraction data is used, the rates of switching may be the only parameters that can be estimated accurately. On the other hand, using cell number data enables accurate estimation of the net division rate for each phenotype, and it can even enable estimation of the state-dependent rates of cell division and cell death. We conclude by applying our framework to a publicly available dataset.

## 1 Introduction

Cancer evolution has long been understood to be a genetic process. However, recent evidence suggests an equally important role for non-genetic forces, including epigenetic mechanisms and the inherent stochasticity in gene transcription and translation [1, 2, 3, 4, 5, 6]. These mechanisms are heritable and reversible, and they can enable cells to dynamically switch between two or more phenotypic states. Such switching dynamics have been observed e.g. in lung cancer [7, 8, 9], melanoma [10, 11, 12], glioblastoma [13, 14], leukemia [15, 16], colon cancer [17, 18, 19, 20] and breast cancer [21, 22, 23, 24]. The different phenotypes commonly show differential responses to drug treatments, which enhances the adaptability of the cancer under treatment and significantly increases the probability of treatment resistance [25].

Unraveling how the cancer-specific rates of cell division, cell death and phenotypic switching shape tumor evolution over time is crucial to furthering our understanding of the disease and to informing new treatment strategies. For example, in a two-phenotype cancer where one type is drug-sensitive and the other is drug-tolerant, the change in phenotypic proportions during the initial stages of treatment can be explained by a combination of sensitive cells dying, drug tolerant cells proliferating, and cells switching between sensitivity and tolerance. Disentangling the relative rates at which these events occur can help us to better understand how resistance arises, how it evolves over time, and how best to combat it [25].

Our current quantitative understanding of the rates of cell proliferation and phenotypic switching in cancer is largely derived from cell line experiments. In these experiments, live cells are commonly sorted into phenotypes, e.g. based on gene expression profiles or cell morphologies, isolated subpopulations are expanded in culture, and phenotypic proportions are tracked over time (Fig. 1). These isolated subpopulations have been observed to give rise to all other phenotypes over time, and to eventually reconstitute a stable distribution of phenotypes that is characteristic of the parental population [21, 17, 19, 20, 23, 24, 14].

**Figure 1:**
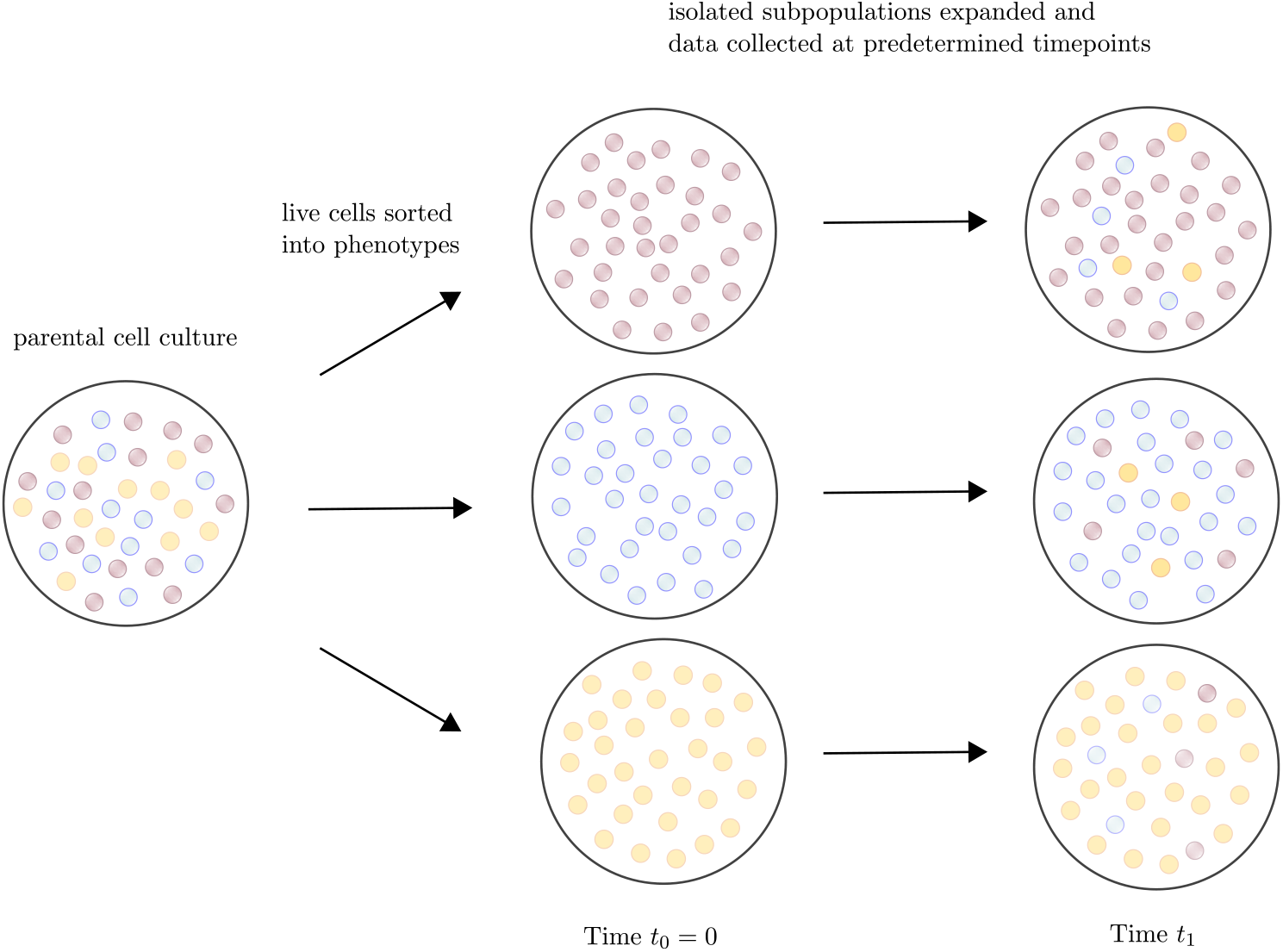
The dynamics of phenotypic switching are commonly interrogated by sorting live cells into isolated phenotypic subpopulations and expanding these subpopulations in culture [21, 17, 19, 20, 23, 24, 14]. By tracking the evolution of phenotypic proportions over time and applying mathematical models of phenotypic switching, it becomes possible to estimate the quantitative parameters of the process [21, 26, 27, 28, 29, 22, 12, 30].

To explain this behavior, simple mathematical models of phenotypic switching have been proposed, and these models have been used to estimate the rates at which cells switch between states [21, 26, 27, 28, 29, 22, 12, 30]. These works are reviewed in Appendix A. Previous estimation methods have been deterministic in nature, and they have generally derived their estimates from data on the fraction of cells in each state at each time point. If the total size of the cell population is measured at the same time points, as e.g. in [30], one obtains data on the number of cells in each state at each time point. We will show that when cell fraction data is used, the rates of phenotypic switching may be the only parameters that can be estimated accurately. In contrast, using cell number data enables accurate estimation of the net cell division rate for each phenotype, and it can even enable estimation of the state-dependent rates of cell division and cell death. Understanding how growth rates vary between types is as important as understanding the rates of phenotypic switching, especially in the context of treatment response. Not only do the growth rates influence the phenotypic composition of the population, they also control the evolution of the tumor burden over time.

Our goal in this work is to develop a statistically rigorous framework for estimating the rates of cell proliferation and phenotypic switching in cancer. In contrast to previous approaches, our framework explicitly models the stochastic dynamics of cell division, cell death and phenotypic switching, it provides likelihood-based confidence intervals for the model parameters, and it enables estimation both from cell fraction and cell number data. We also use our framework to analyze the identifiability of model parameters and how it depends on the input data. This important topic has not been addressed by previous works. The rest of the paper is organized as follows. In Section 2, we introduce our stochastic model of cell division, cell death and phenotypic switching. In Section 3, we state our assumptions on the cell line experiments conducted and the data collected. In Sections 4 and 5, we propose statistical models for cell number and cell fraction data, respectively, and show how parameter estimates and confidence intervals are computed. In Section 6, we present theoretical analysis of the identifiability of parameters under each model. In Section 7, we conduct numerical experiments, and in Section 8, we apply our framework to a publicly available dataset. We conclude with a discussion section (Section 9).

## 2 Multitype branching process model

### 2.1 Model definition and model parameters

To model the cell population dynamics, we employ a multitype branching process model in continuous time, with *K* ≥ 2 types [31]. In the model, a type-*j* cell divides into two cells at rate *b*_*j*_ ≥ 0, it dies at rate *d*_*j*_ ≥ 0, and it switches to type-*k* at rate *ν*_*jk*_ ≥ 0 for *k* ≠ *j*, independently of all other cells. This means that in an infinitesimally short time interval of length Δ*t >* 0, a type-*j* cell divides with probability *b*_*j*_Δ*t*, it dies with probability *d*_*j*_Δ*t*, and it switches to type *k* with probability *ν*_*jk*_Δ*t*. The multitype branching process model captures a variety of switching dynamics previously observed in the literature (Fig. 2).

**Figure 2:**
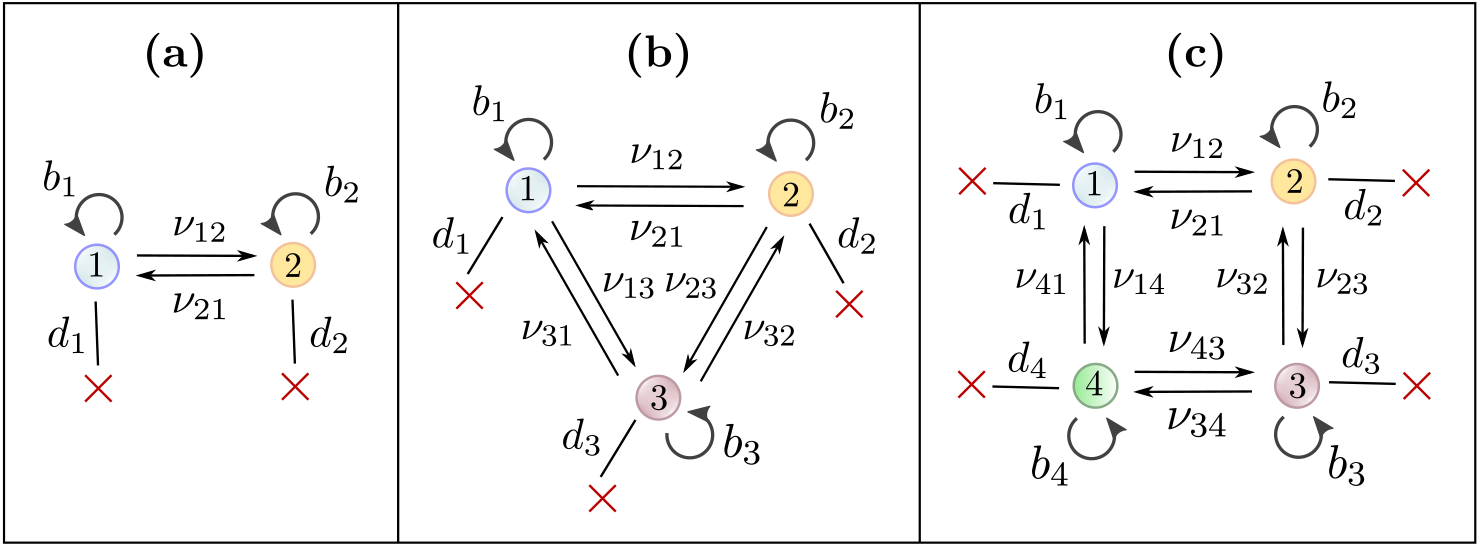
The multitype branching process model captures a variety of switching dynamics previously observed in the literature. **(a)** A two-type model captures e.g. the dynamics between HER2+ and HER2− cell states in Brx-82 and Brx-142 breast cancer cells [23]. **(b)** A three-type model captures e.g. the dynamics between stem-like, basal and luminal cell states in SUM149 and SUM159 breast cancer cells [21]. **(c)** A four-type model captures e.g. the dynamics between CD24^Low^/ALDH^High^, CD24^Low^/ALDH^Low^, CD24^High^/ALDH^High^ and CD24^Hich^/ALDH^Low^ cell states in GBC02, SCC029B and SCC070 oral cancer cells [32].

We allow *ν*_*jk*_ = 0 for some *j* and *k*, which means that a type-*j* cell is not able to switch directly to type-*k*. However, in our exposition, we assume that the model is *irreducible*, in that each cell type is accessible from any other cell type, possibly through intermediate types. In other words, for each *j, k* = 1, …, *K* with *k* ≠ *j*, we assume that there exist *r* ≥ 0 integers *m*_1_, …, *m*_*r*_ ∈ *{*1, …, *K*} so that 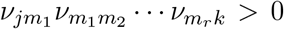. The reducible case is treated in Appendix B below.

For *j* = 1, …, *K*, we define *λ*_*j*_ := *b*_*j*_ − *d*_*j*_ as the net birth rate of a type-*j* cell. We collect the growth parameters into 1 *× K* vectors **b** = (*b*_1_, …, *b*_*K*_), **d** = (*d*_1_, …, *d*_*K*_) and ***λ*** = (*λ*_1_, …, *λ*_*K*_). We collect the switching rates into a *K ×* (*K* − 1) matrix ***ν***, where the *j*-th row vector is (*ν*_*jk*_)_*k j*_. We finally define ***λ***^[−*j*]^ := ***λ*** − *λ*_*j*_**1** as the vector of net birth rates relative to *λ*_*j*_, with 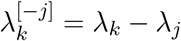 for *k* ≠ *j* and 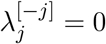.

### 2.2 Stochastic processes

Let **n** = (*n*_1_, …, *n*_*K*_) be the 1*×K* vector of starting cell numbers of each type. The state of the model at time *t* ≥ 0 is encoded in the 1*×K* vector of cell numbers 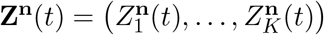, where 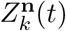 is the number of type-*k* cells at time *t*. We will commonly deal with the case **n** = **e**_*j*_ for some *j* = 1, …, *K*, where **e**_*j*_ is the *j*-th unit vector, meaning that the process is started by a single type-*j* cell. We write 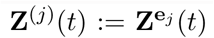 for that case, and we let **m**^(*j*)^(*t*) and **Σ**^(*j*)^(*t*) denote the mean vector and covariance matrix for **Z**^(*j*)^(*t*):

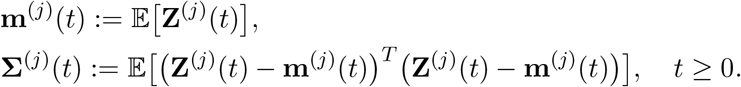

On the event 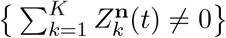, we let **Δ**^**n**^(*t*) denote the vector of cell fractions, i.e.

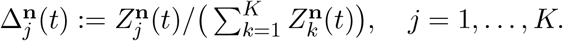

As above, we write 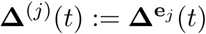 for *j* = 1, …, *K*.

### 2.3 Infinitesimal generator and mean matrix

We define the *K × K* matrix **A** with *a*_*jj*_ := *λ*_*j*_ Σ _*k*≠*j*_ *ν*_*jk*_ for *j* = 1, …, *K* and *a*_*jk*_ := *ν*_*jk*_ for *k* ≠ *j* as the *infinitesimal generator* of the model, where *a*_*jk*_ is the net rate at which a cell of type *j* produces a cell of type *k*. We next define the matrix exponential

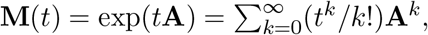

which is referred to as the *mean matrix*. It can be shown that the *j*-th row vector of **M**(*t*) is **m**^(*j*)^(*t*) = 𝔼 [**Z**^(*j*)^(*t*)], i.e. the mean vector for the process started by a single type-*j* cell [31]. Note that **A** and **M**(*t*) depend only on the switching rates ***ν*** and the net birth rates ***λ***.

### 2.4 Long-run behavior

In the absence of phenotypic switching, the subpopulation of type-*j* cells grows at exponential rate *λ*_*j*_ on average. However, in the presence of phenotypic switching, and under the assumption of irreducible switching dynamics, all subpopulations eventually grow at the same rate *σ*. More precisely, there exists a real number *σ* and positive 1 *× K* vectors ***β*** = (*β*_1_, …, *β*_*K*_) and ***γ*** = (*γ*_1_, …, *γ*_*K*_) so that

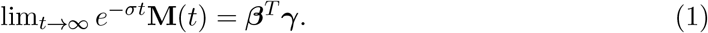

This means that if the process is started by a single type-*j* cell, the mean number of type-*k* cells at time *t* is approximately *β*_*j*_*γ*_*k*_*e*^*σt*^ when *t* is large. It follows that if we define

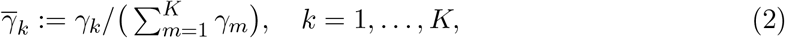

then 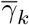 is the long-run proportion of type-*k* cells in the population, independently of the initial condition. Thus, in the long run, cell proportions tend towards an equilibrium distribution given by 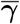, which is consistent with the experimental observations discussed in the introduction. For the mathematical details, including how to compute ***β*** and ***γ***, see e.g. Section V.7 of [31].

## 3 Experiments and data

We assume that each experiment is started with a known initial condition, encoded by the 1 *× K* vector **n** = (*n*_1_, …, *n*_*K*_) of starting cell numbers for each type. We let *I* ≥ 1 denote the number of distinct initial conditions and **n**_*i*_ = (*n*_*i*1_, …, *n*_*iK*_) denote the *i*-th initial condition. We assume that for each *i* = 1, …, *I* and *j* = 1, …, *K*, either *n*_*ij*_ = 0 or *n*_*ij*_ is large, which is generally the case for the cell line experiments discussed in the introduction.

We define 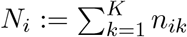 as the total number of starting cells in the *i*-th condition and **f**_*i*_ = (*f*_*i*1_, …, *f*_*iK*_) as the vector of starting cell fractions, with *f*_*ij*_ := *n*_*ij*_*/N*_*i*_. We let *L* ≥ 1 be the number of time points at which data is collected and we let 0 *< t*_1_ *< t*_2_ *< … < t*_*L*_ denote the time points. Finally, we let *R* ≥ 1 be the number of experimental replicates performed. In the development of our estimation framework, we assume that each experiment returns measurements from a single time point only, meaning that the sample is discarded once measurements are taken (endpoint data). This is e.g. the case when fluorescence-activated cell sorting (FACS) is used to identify phenotypes. Sometimes, the data collected is sequential, meaning that a single experiment returns measurements from multiple time points. This is e.g. the case when live cell imaging is used. While our framework is developed for endpoint data, we show in Section 7 that it also yields reasonable estimates for sequential data.

The data collected in each experiment is either a vector **n**_*i,ℓ,r*_ = (*n*_*i,ℓ,r*,1_, …, *n*_*i,ℓ,r,K*_) of cell numbers or **f**_*i,ℓ,r*_ = (*f*_*i,ℓ,r*,1_, …, *f*_*i,ℓ,r,K*_) of cell fractions. Here, *n*_*i*,, *r,j*_ is the number of type-*j* cells in the *r*-th replicate of the experiment started by the *i*-th initial condition and ended at the *ℓ*-th timepoint, and *f*_*i,ℓ,r,j*_ is the corresponding cell fraction.

## 4 Estimation for cell number data

In this section, we propose a maximum likelihood estimation framework for cell number data. Our framework is rooted in a central limit theorem for the vector **Z**^**n**^(*t*) of cell numbers at time *t*, which approximates the distribution of **Z**^**n**^(*t*) by a normal distribution for a large starting population. Using the central limit theorem, we propose a statistical model and derive a likelihood function for cell number data, which is then used to compute maximum likelihood estimates and confidence intervals for all parameters.

### 4.1 Central limit theorem

We begin by establishing a central limit theorem (CLT) for the vector **Z**^**n**^(*t*) of cell numbers, which involves decomposing the branching process (**Z**^**n**^(*s*))_*s*≥0_ into i.i.d. processes started by single cells. By fixing the vector of starting cell proportions, and sending the starting population size to infinity, we can apply the standard (multivariate) CLT to obtain the following result.

#### Proposition 1.

*Let* ***α*** *be* 1 *× K with α*_*i*_ ≥ 0 *for i* = 1, …, *K and* 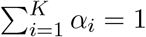. *Define*

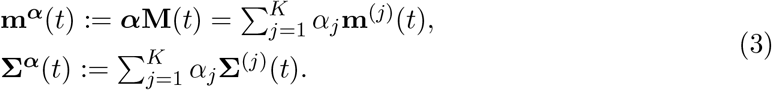

*Let J* ≥ 1 *be any integer. For any K × J matrix* **C**, *then as N* → ∞,

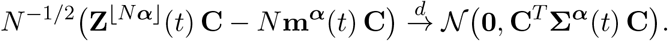

*Here, the covariance matrix* **Σ**^(*j*)^(*t*) *is given by*

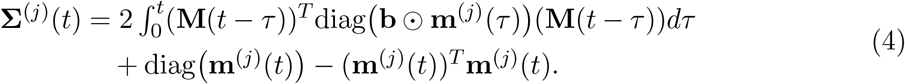

*Proof*. Appendix E.

Note that in Proposition 1, some coordinates of the vector ***α*** of starting cell proportions are allowed to be 0. In the *N* → ∞ regime, the starting condition ⌊*N* ***α*** ⌋ will therefore either include no cell or a large number of cells of any given type. This is consistent with our assumptions on the vectors **n**_1_, …, **n**_*I*_ of experimental starting conditions (Section 3).

Also note that Proposition 1 is established for linear transformations **Z** ^**⌊**N**α⌋**^ (*t*) **C** of **Z** ^**⌊**N**α⌋**^ (*t*). This allows us to obtain a CLT for cases where we do not observe the full vector **Z** ^*N****α***^ (*t*). For example, if we set **C** := **1**^*T*^, then 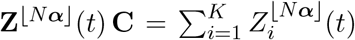 is the total number of cells at time *t*. This more general CLT also becomes useful when estimating from models with reducible switching dynamics, as we discuss in Appendix B.

### 4.2 Statistical model

Based on Proposition 1, we propose the following statistical model for the data **n**_*i,ℓ,r*_:

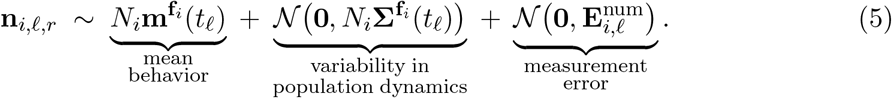

The first two terms capture the mean and variance of the population dynamics, in accordance with Proposition 1, while the final term captures experimental measurement error.

The vectors **n**_*i*,, *r*_ and **n**_*j,m,s*_ are assumed independent for (*i, l, r*) ≠ (*j, m, s*), and they are assumed i.i.d. for (*i, l*) = (*j, m*) and *r* ≠ *s*. This assumes that data from distinct time points come from distinct experiments (endpoint data). We make this assumption since the CLT of Proposition 1 holds for the distribution of **Z** ^⌊N*α*⌋^ (*t*) at a fixed time point *t*. Developing an analogous statistical model for sequential data requires extending the CLT in Proposition 1 to a process-level or functional CLT. As stated above, we will show in Section 7 that the statistical model in (5) yields reasonable estimates for sequential data.

Note that the mean behavior 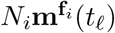 of the model depends only on ***ν*** and ***λ***, while the variance term 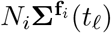 depends on ***ν, λ*** and **b** by Proposition 1. It is therefore natural to parametrize the first two terms in (5) by **b, *λ, ν*** instead of the primary parameters **b, d, *ν***. We assume that the *K × K* covariance matrix 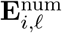 associated with measurement error can be written as a function of **b, *λ, ν*** and additional error parameters 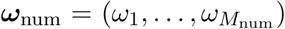 for some *M*_num_ ≥ 0. A simple example is 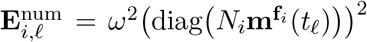 for some *ω >* 0, where the measurement error is assumed to be uncorrelated between types, and to scale with the mean experimental outcomes. We let ***θ***_num_ = [**b, *λ, ν, ω***_num_] be the complete 1 *× K*(*K* + 1) + *M*_num_ vector of model parameters including the error parameters.

### 4.3 Maximum likelihood estimates and confidence intervals

From the statistical model (5), it is straightforward to derive a likelihood function, i.e. the probability of observing the experimental data as a function of the model parameters:

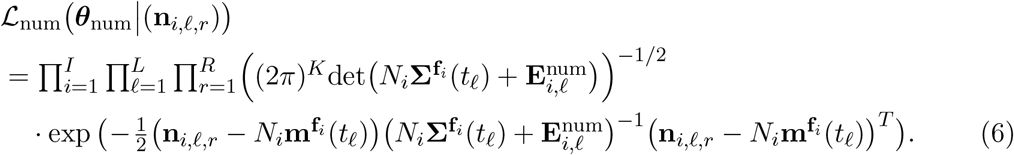

We next define the negative double log-likelihood,

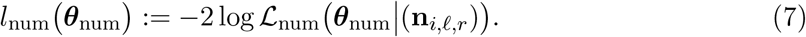

The maximum likelihood estimate 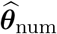 for the parameter vector ***θ***_num_ is obtained by minimizing *ℓ*_num_(***θ***_num_) over a set of feasible parameters **Θ**_num_:

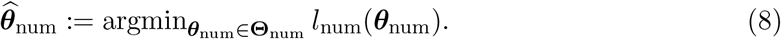

In the feasible set **Θ**_num_, we restrict the parameter values so that ***ν*** ≥ **0, b** ≥ **0** and ***λ*** ≤ **b**.

Further restrictions can be made depending on the context, see e.g. Appendix B.

A 1 − *α* likelihood-based confidence interval 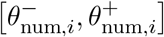 for the *i*-th model parameter *θ*_num,*i*_ can be obtained by collecting all values *θ* for which the null hypothesis *θ*_num,*i*_ = *θ* is accepted under a likelihood ratio test. The endpoints 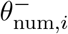 and 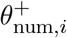 can be computed by solving the following two constrained optimization problems:

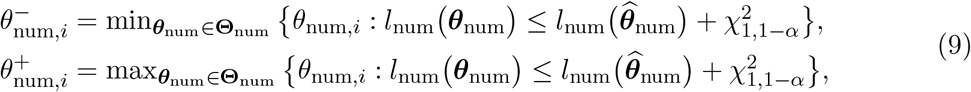

where 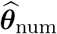 is the MLE estimator defined by (8) and 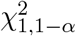 is the (1 − *α*)-th quantile of the 𝒳^2^-distribution. See e.g. [33,34,35,36,37] for a further discussion.

Our estimation framework is based on solving the optimization problems in (8) and (9) using the sqp solver in MATLAB.

## 5 Estimation for cell fraction data

In this section, we propose a maximum likelihood estimation framework for cell fraction data. As for cell number data, the framework is rooted in a central limit theorem for the vector of cell fractions **Δ**^**n**^(*t*), which we use to propose a statistical model and derive a likelihood function for cell fraction data, and to then compute maximum likelihood estimates and confidence intervals for the model parameters.

### 5.1 Central limit theorem

We begin by establishing a central limit theorem for the vector of cell fractions **Δ**^**n**^(*t*). This CLT has already been established for the case of an isolated large starting population by Yakovlev and Yanev [38]. We extend their argument to more general starting conditions by fixing the vector ***α*** of starting cell proportions and sending the total population size *N* to infinity. We also provide a simplified expression for the covariance matrix **S**^***α***^(*t*) and show that the mean function **p**^***α***^(*t*) can be written solely in terms of ***ν*** and ***λ***^[−*j*]^ for any *j* = 1, …, *K*.

#### Proposition 2.

*Let* ***α*** *be* 1 *× K with α*_*i*_ ≥ 0 *for i* = 1, …, *K and* 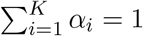. *Define*

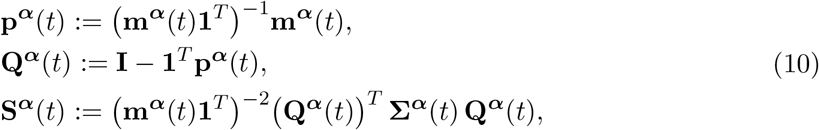

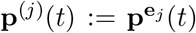 *and* 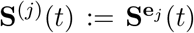, *where* **m**^***α***^(*t*) *and* **Σ**^***α***^(*t*) *are defined as in* (3). *Let J* ≥ 1 *be any integer. For any K × J matrix* **C**, *then as N* → ∞,

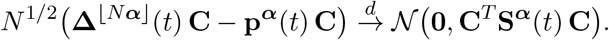

*Here, the mean function* **p**^***α***^(*t*) *can be written solely as a function of the switching rates* ***ν*** *and the relative net birth rates* ***λ***^[−*j*]^ *for any j* = 1, …, *K*.

*Proof*. Appendix F.

### 5.2 Statistical model and estimation

Based on Proposition 2, we propose the following statistical model for the data **f**_*i,ℓ,r*_:

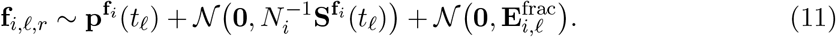

Note that the mean behavior 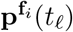 depends only on ***ν*** and ***λ***^[−1]^, while the variance term 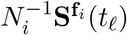 depends on all model parameters ***ν, λ***^[−1]^, *λ*_1_, **d**. The choice of type-1 as a reference phenotype is arbitrary, and we use **d** as opposed to **b** as we found it to perform well numerically. We assume that the *K × K* covariance matrix 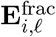 associated with measurement error can be written as a function of **d**, *λ*_1_, ***λ***^[−1]^, ***ν*** and added error parameters 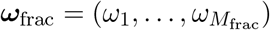 for some *M*_frac_ ≥ 0. A simple example is 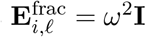 for some *ω >* 0, where the measurement error is assumed uncorrelated between cell types, and of the same magnitude for all initial conditions and all time points. We let ***θ***_frac_ = [**d**, *λ*_1_, ***λ***^[−1]^, ***ν, ω***_frac_] denote the complete vector of model parameters including the error parameters.

When deriving a likelihood function for the statistical model (11), we note that the last coordinate of **f**_*i*,, *r*_ provides no new information over the first *K* − 1 coordinates, since the coordinates always sum to one. In the likelihood function, we therefore only consider the first *K* − 1 coordinates, which we can accomplish by multiplying **f**_*i,ℓ,r*_ by the *K ×* (*K* − 1) matrix **B** with 1 on the diagonal and 0 off it. In this way, we obtain the following likelihood:

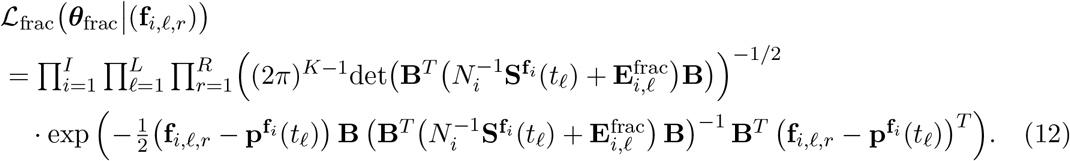

As for cell number data, we define the negative double log-likelihood,

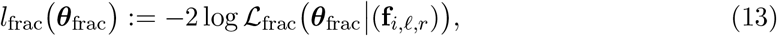

and obtain the maximum likelihood estimate for ***θ***_frac_ by solving

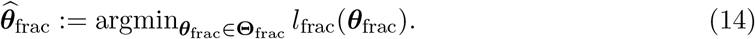

In the feasible set **Θ**_frac_, we restrict the parameter values so that ***ν*** ≥ **0, d** ≥ **0**, *λ*_1_ ≥ −*d*_1_ and (*λ*_*j*_ − *λ*_1_) + *d*_*j*_ + *λ*_1_ ≥ 0 for *j* = 2, …, *K*. Further restrictions can be made depending on the context, see e.g. Section 8 and Appendix C.

The computation of confidence intervals proceeds as described in Section 4.3.

## 6 Structural identifiability

In this section, we analyze the structural identifiability of the statistical models of Sections 4 and 5. Informally, structural identifiability refers to whether a parameter can be estimated accurately given an infinite amount of noise-free data. More precisely, a parameter is structurally identifiable if complete knowledge of the model distribution uniquely determines the value of the parameter, in the absence of any measurement error [39, 40].

Here, we will assume that we know the behavior of the mean functions **m**^(*j*)^(*t*) and **p**^(*j*)^(*t*) and the covariance functions **Σ**^(*j*)^(*t*) and **S**^(*j*)^(*t*) close to time 0, and we will analyze to what extent the model parameters can be extracted from this information. In other words, we are interested in the following question: If we conduct experiments started from isolated subpopulations, and perfect observations are made of the first two statistical moments of the model close to time 0, can we identify the model parameters?

The analysis in this section serves two purposes. First of all, it ascertains whether in this idealized setting, the model parameters can be extracted uniquely from short-term observations of the population dynamics. Second, the analysis indicates how much information is required to estimate each model parameter accurately, which yields valuable insights into how comparatively difficult it is to estimate the parameters from more limited data.

### 6.1 Cell number data

In the following proposition, we show that for cell number data, the switching rates ***ν*** and the net birth rates ***λ*** can be recovered uniquely from knowledge of the mean functions **m**^(*j*)^(*t*) close to time 0, while the birth rates **b** can be recovered from the covariance matrices **Σ**^(*j*)^(*t*).

#### Proposition 3.

1. *For each j* = 1, …, *K, the s*I*witching rates ν*_*jk*_, *k* ≠ *j, and the net birth rate λ*_*j*_ *are uniquely determined by* 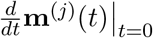
2. *For each j* = 1, …, *K, if the switching rates ν*_*jk*_, *k* ≠ *j, and the net birth rate λ*_*j*_ *are known, the birth rate b*_*j*_ *is uniquely determined by* 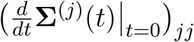.

*Proof*. Appendix G.

Proposition 3 establishes the structural identifiability of all model parameters for cell number data. The process of extracting the parameters as suggested by Proposition 3 can be thought of as follows: If we want to know *ν*_jk_ for some *k* … *j*, we can simply plot the mean function 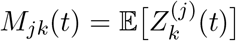 and compute its slope at 0. If we want to know the birth rate *b*_*j*_, we can plot the variance function 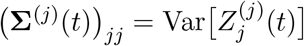 and compute its slope at 0.

It is important to note that we are not suggesting to use this approach to estimate parameters from real data. Instead, we are establishing theoretically that there is sufficient information in the distribution of the data close to time 0 to determine all model parameters uniquely. In particular, we can in theory predict the entire evolutionary trajectory of the population from short-term observations of the initial population dynamics.

### 6.2 Cell fraction data

In the following proposition, we show that for cell fraction data, only the switching rates ***ν*** can be recovered from the slopes of the mean functions **p**^(*j*)^(*t*) at time 0. The net birth rate differences ***λ***^[−1]^ can be recovered from the curvatures of the mean functions at time 0 or from the equilibrium proportions 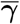 between cell types if they are known. We are not able to learn any more parameters from the mean functions, since **p**^(*j*)^(*t*) can be written solely as a function of ***ν*** and ***λ***^[−1]^ by Proposition 2. The slopes of the covariance functions **S**^(*j*)^(*t*) depend only on ***ν***, meaning that they provide no extra information on the model parameters.

#### Proposition 4.

1. *F*I*or j* = 1, …, *K, the switching rates ν*_*jk*_, *k* ≠ *j, are uniquely determined by* 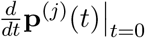.
2. *If the switching rates* ***ν*** *are known, the net birth rate differences* ***λ***^[−1]^ *are uniquely determined by (i)* 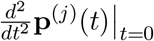 *for j* = 1, …, *K or (ii) the equilibrium proportions* 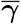.
3. *For* 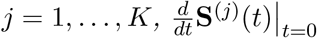 *only depends on the switching rates ν*_*jk*_ *for k* ≠ *j*.

*Proof*. Appendix H.

As for the remaining model parameters, the net birth rate *λ*_1_ and the birth rates **b**, they require information on the curvatures of the covariance functions **S**^(*j*)^(*t*) at time 0 at the least. We will not analyze the structural identifiability of these parameters further. Proposition 4 indicates that one should not expect to be able to estimate these parameters accurately from cell fraction data, which is confirmed by numerical experiments in Section 7.3.

### 6.3 Comparison

The results of our identifiability analysis are summarized in Table 2. Our analysis indicates that the switching rates ***ν*** and net birth rates ***λ*** are easy to estimate for cell number data, using information only on the mean behavior of the population. The birth rates **b** are harder to estimate, since they require second moment information, but they may still be obtainable with sufficient data, as we discuss further in Section 7. For cell fraction data, the switching rates ***ν*** are easy to estimate using the mean behavior of the population. The net birth rate differences ***λ***^[−1]^ can also be estimated from the mean, but they require more information. The remaining model parameters are unlikely to be obtainable from real datasets.

**Table 1:** Notation for the stochastic model of Section 2.

| Symbol | Dimension | Description |
| --- | --- | --- |
| $K$ | 1 | Number of types |
| $b_j$ | 1 | Division rate of type-$j$ cells |
| $d_j$ | 1 | Death rate of type-$j$ cells |
| $\nu_{jk}$ | 1 | Rate of switching from type-$j$ to type-$k$ |
| $\lambda_j$ | 1 | Net birth rate of type-$j$ cells, $\lambda_j = b_j - d_j$ |
| $\lambda_k^{[-j]}$ | 1 | Net birth rate relative to $\lambda_j$, $\lambda_k^{[-j]} = \lambda_k - \lambda_j$ |
| $\mathbf{A}$ | $K \times K$ | Infinitesimal generator of the branching process model |
| $\mathbf{M}(t)$ | $K \times K$ | Mean matrix at time $t$, $\mathbf{M}(t) = \exp(t\mathbf{A})$ |
| $\bar{\gamma}$ | $1 \times K$ | equilibrium proportions between cell types |
| $\mathbf{Z}^{\mathbf{n}}(t)$ | $1 \times K$ | Vector of cell numbers at time $t$, initial condition $\mathbf{n}$ |
| $\mathbf{Z}^{(j)}(t)$ | $1 \times K$ | Vector of cell numbers at time $t$, initial condition $\mathbf{e}_j$ |
| $\mathbf{m}^{\alpha}(t)$ | $1 \times K$ | $\mathbf{m}^{\alpha}(t) = \alpha \mathbf{M}(t) = \sum_{j=1}^K \alpha_j \mathbf{m}^{(j)}(t)$ |
| $\mathbf{m}^{(j)}(t)$ | $1 \times K$ | $\mathbf{m}^{(j)}(t) = \mathbb{E}[\mathbf{Z}^{(j)}(t)] = \mathbf{e}_j \mathbf{M}(t)$ |
| $\Sigma^{\alpha}(t)$ | $K \times K$ | $\Sigma^{\alpha}(t) = \sum_{j=1}^K \alpha_j \Sigma^{(j)}(t)$ |
| $\Sigma^{(j)}(t)$ | $K \times K$ | $\Sigma^{(j)}(t) = \mathbb{E}[(\mathbf{Z}^{(j)}(t) - \mathbf{m}^{(j)}(t))^T (\mathbf{Z}^{(j)}(t) - \mathbf{m}^{(j)}(t))]$ |
| $\Delta^{\mathbf{n}}(t)$ | $1 \times K$ | Vector of cell fractions at time $t$, initial condition $\mathbf{n}$ |
| $\mathbf{p}^{\alpha}(t)$ | $1 \times K$ | $\mathbf{p}^{\alpha}(t) = (\alpha \mathbf{M}(t) \mathbf{1}^T)^{-1} \alpha \mathbf{M}(t)$ |
| $\mathbf{Q}^{\alpha}(t)$ | $K \times K$ | $\mathbf{Q}^{\alpha}(t) = \mathbf{I} - \mathbf{1}^T \mathbf{p}^{\alpha}(t)$ |
| $\mathbf{S}^{\alpha}(t)$ | $K \times K$ | $\mathbf{S}^{\alpha}(t) = (\mathbf{m}^{\alpha}(t) \mathbf{1}^T)^{-2} (\mathbf{Q}^{\alpha}(t))^T \Sigma^{\alpha}(t) \mathbf{Q}^{\alpha}(t)$ |

**Table 2:** Table 2 Summary of the structural identifiability analysis of Propositions 3 and 4. For cell number data, the switching rates ***ν*** and the net birth rates ***λ*** are identifiable from the slopes (first derivatives) of the mean functions **m**^(*j*)^(*t*) (first moments) at time 0. The birth rates **b** are identifiable from the slopes of the covariance functions **Σ**^(*j*)^(*t*) (second moments). For cell fraction data, only the switching rates ***ν*** are identifiable from the slopes of the mean functions **p**^(*j*)^(*t*), while the net birth rate differences ***λ***^[−1]^ can be determined from their curvatures (second derivatives). In contrast to cell number data, the slopes of the covariance functions **S**^(*j*)^(*t*) for cell fraction data provide no extra information on the model parameters.

| Moment | Derivative | Cell number data | Cell fraction data |
| --- | --- | --- | --- |
| 1 | 1 | $\boldsymbol{\nu}, \boldsymbol{\lambda}$ | $\boldsymbol{\nu}$ |
| | 2 | — | $\boldsymbol{\lambda}^{[-1]}$ |
| 2 | 1 | $\mathbf{b}$ | $\boldsymbol{\nu}$ |

## 7 Numerical experiments

In this section, we apply our maximum likelihood framework to computer-generated data, and we explore how well the model parameters can be estimated depending on what data is collected. In all cases, we assume that experiments are conducted from isolated initial conditions, and we assume no measurement noise, i.e. 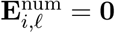 and 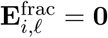.

### 7.1 Implementation in MATLAB

Our estimation framework has been implemented in MATLAB codes which are available at https://github.com/egunnars/phenotypic_switching_inference/. The framework returns (i) a maximum likelihood estimate and (ii) a likelihood-based confidence interval for each parameter, using the sequential quadratic programming (sqp) solver in MATLAB. Before solving the maximum likelihood problem, we compute initial parameter estimates from simpler models, which we use to initialize the optimization and to rescale the model parameters so that they are of similar magnitude. In most cases, we have found it sufficient to solve the maximum likelihood problem once, starting from the simple estimates. However, our MATLAB codes provide the option to solve the problem several times using different initial guesses. Details of the implementation are provided in Appendix C.

### 7.2 Illustrative example

For illustrative purposes, we first show a graphical depiction of the output of our estimation framework for a single dataset. We generated artificial cell number and cell fraction data by performing a stochastic simulation of the branching process model from Section 2. We then used the data to compute MLE estimates and confidence intervals for the model parameters. The data was generated assuming *K* = 2 cell types, *I* = 2 isolated initial conditions, *L* = 6 time points, and *R* = 3 replicates. Estimation results are shown in Figure 3.

**Figure 3:**
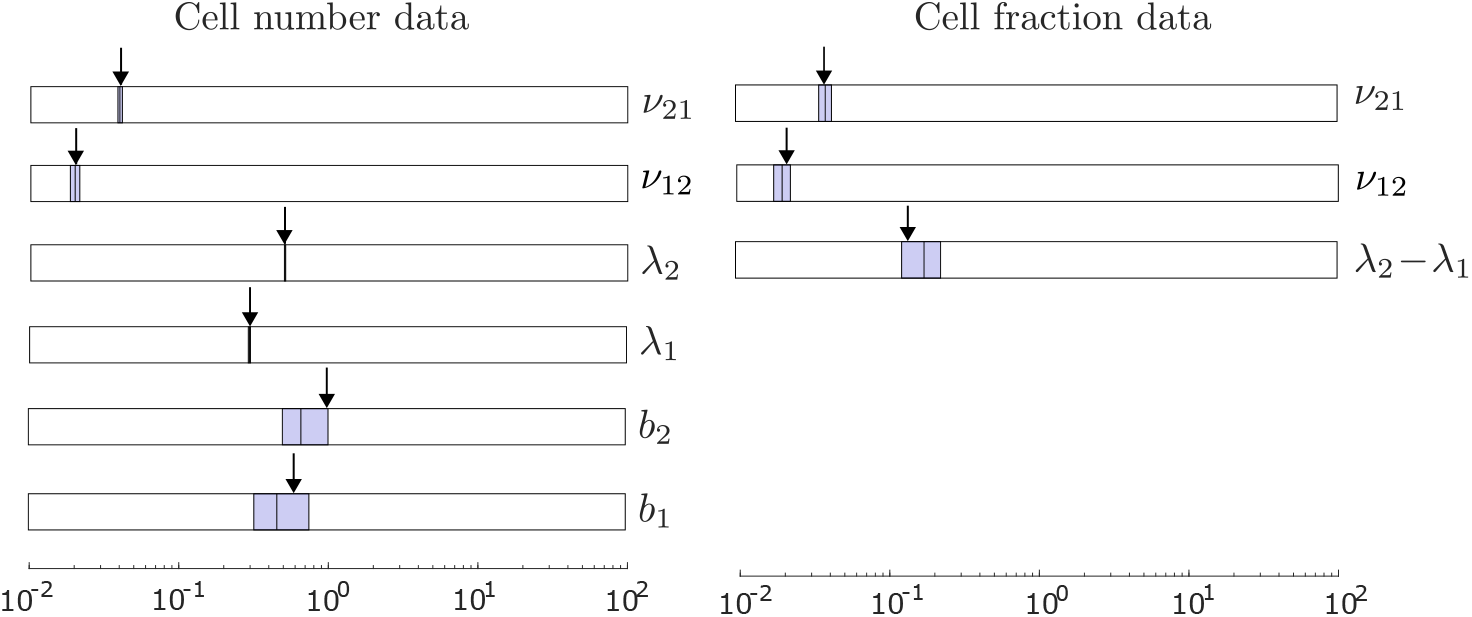
Graphical depiction of the output of our estimation framework. We first generated artificial cell-number and cell-fraction data by simulating the branching process model of Section 2 for *b*_1_ = 0.6, *d*_1_ = 0.3, *b*_2_ = 1.0, *d*_2_ = 0.5, *ν*_12_ = 0.02, *ν*_21_ = 0.04, **f**_1_ = [1, 0], **f**_2_ = [0, 1] and *N*_1_ = *N*_2_ = 1, 000. Using this data, we computed maximum likelihood estimates and likelihood-based 95% confidence intervals (CIs) for the model parameters. For each parameter, the shaded region indicates the CI, the vertical bar inside the interval indicates the MLE estimate, and the arrow points to the true value of the parameter.

Note first the difference in scale between the switching rates and the rates involving cell division and death. This is typically the case, since epigenetic modifications can generally be retained for 10–10^5^ cell divisions [41, 3]. Also note that all model parameters are estimated more accurately for cell number data than cell fraction data, in that their confidence intervals are narrower for cell number data. Otherwise, the relative accuracy with which different model parameters can be estimated is in line with our identifiability analysis in Section 6.

### 7.3 Estimation across a wide range of biologically realistic regimes

For a more thorough evaluation of estimation accuracy, we generated 10,000 artificial datasets for *K* = 2 cell types. We first generated 100 biologically realistic parameter regimes and then generated 100 datasets for each regime. To generate the parameter regimes, we sampled birth and death rates uniformly between 0 and 1, and sampled switching rates log-uniformly between 10^−1^ and 10^−3^. We considered both regimes where the two phenotypes have positive net birth rates (*λ*_1_, *λ*_2_ *>* 0) and regimes where one phenotype has a negative net birth rate (*λ*_1_ *<* 0, *λ*_2_ *>* 0). The latter regimes are relevant to the dynamics of anti-cancer treatment response, where one phenotype is drug-sensitive and the other is drug-tolerant. As in Section 7.2, we assumed *I* = 2 isolated initial conditions, *L* = 6 time points and *R* = 3 replicates. Further details of the data generation are provided in Appendix D.

For each dataset, we used our framework to compute MLE estimates for all model parameters. In this way, we obtained 100 estimates of each parameter under each parameter regime, which we used to compute the coefficient of variation (CV) for the MLE estimator of the parameter. The CV is the sample standard deviation of the MLE estimator as a proportion of its sample mean, and it measures the percentage error in the estimation.

The results are shown in Figure 4. A horizontal line is drawn at 25% CV to indicate whether parameters can be estimated with reasonable accuracy. Note that for the switching rates ***ν***, the median CV for cell fraction data is about twice as large as for cell number data.

**Figure 4:**
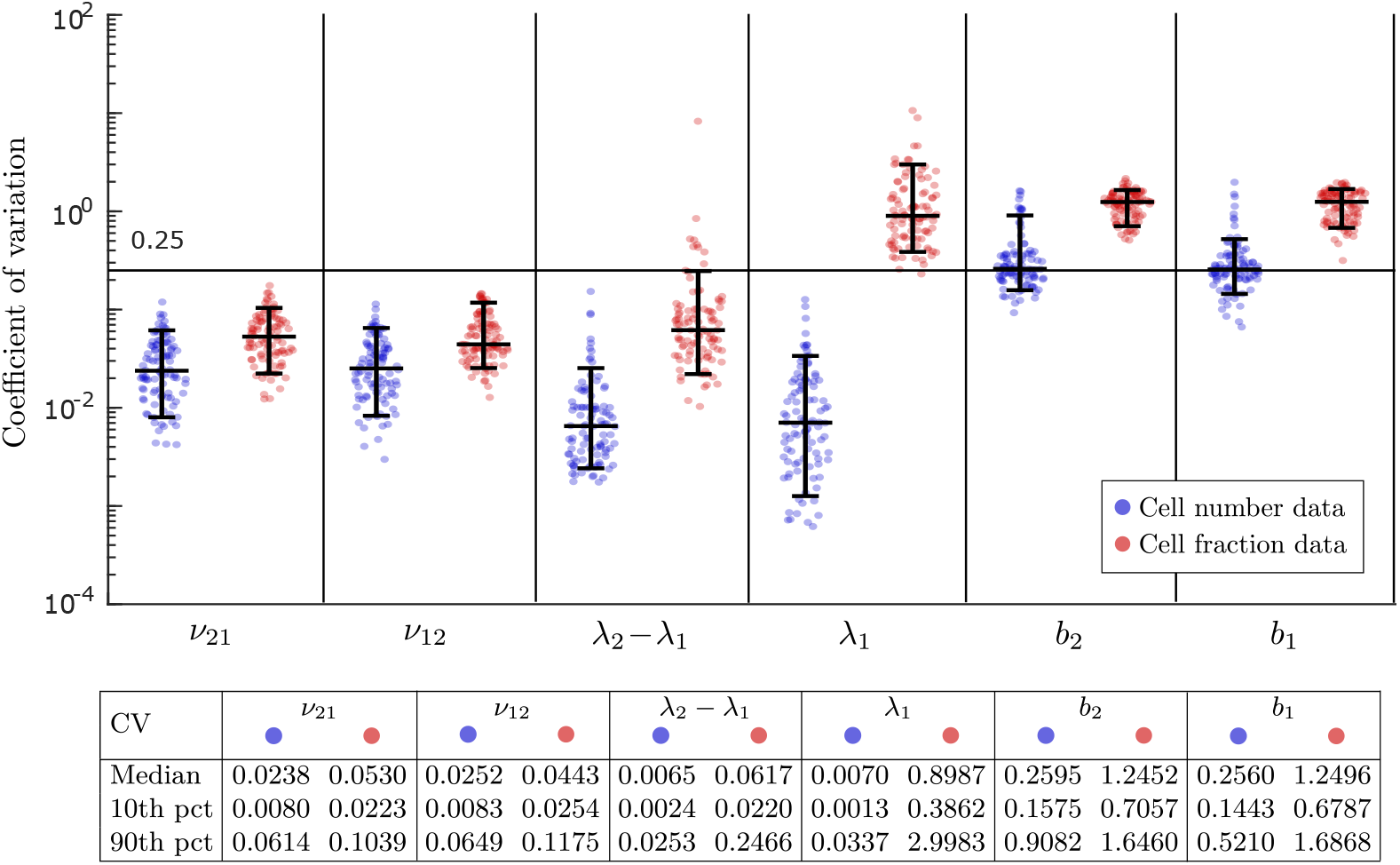
Assessment of estimation error across a wide range of biologically realistic parameter regimes. We first generated 100 different parameter regimes, then generated 100 artifical datasets for each regime, and finally computed parameter estimates for each dataset. To generate the parameter regimes, we sampled birth and death rates uniformly between 0 and 1, and sampled switching rates log-uniformly between 10^−1^ and 10^−3^ (Appendix D). For each parameter and each parameter regime, we used the 100 estimates to compute the coefficient of variation (CV) for the estimates, which measures the error in the estimation. Each dot in the figure represents the CV for a single parameter under a single regime, with the blue dots (resp. red dots) representing estimates from cell number data (resp. cell fraction data). Collectively, the dots enable comparison of estimation error between different model parameters and between cell number and cell fraction data. The horizontal bars represent the 10th percentile, median and 90th percentile of the CVs, bottom to top.

The median CV for the net birth rate difference *λ*_2_ − *λ*_1_ is an order of magnitude larger for cell fraction data than cell number data, and it is two orders of magnitude larger for the net birth rate *λ*_1_. The birth rates **b** can in many cases be estimated reasonably well for cell number data, whereas they are never estimated accurately for cell fraction data. These results are very much in line with our identifiability analysis in Section 6.

Note that for cell fraction data, the estimation error for the net birth rate difference *λ*_2_−*λ*_1_ exceeds the 25% threshold CV for several parameter regimes. This occurs when *λ*_2_ − *λ*_1_ is small in magnitude, more precisely when it is smaller than 0.1 in regimes where the birth rates lie between 0.1 and 1. Note in contrast that for cell number data, the estimation error for *λ*_2_ − *λ*_1_ never exceeds the 25% threshold. This indicates that for cell fraction data, it may be difficult to distinguish the net birth rate difference *λ*_2_ − *λ*_1_ from 0 unless it is relatively pronounced. We discuss this point further in Section 8 below.

### 7.4 Experimental design: Adding replicates vs. adding time points

In this section, we discuss how our framework can be used to evaluate to what extent additional data can improve parameter estimates and to identify experimental designs that best accomplish this goal. To illustrate this point, we compared the effect of (i) doubling the number of replicates from *R* = 3 to *R* = 6 (design 1), (ii) doubling the number of time points from *T* = 6 to *T* = 12, adding time points in between the previous time points (design 2), and (iii) doubling the number of time points, adding time points after the previous points (design 3) (Appendix D). We generated 10 parameter regimes and 100 datasets for each regime. The results are shown in Figure 5.

**Figure 5:**
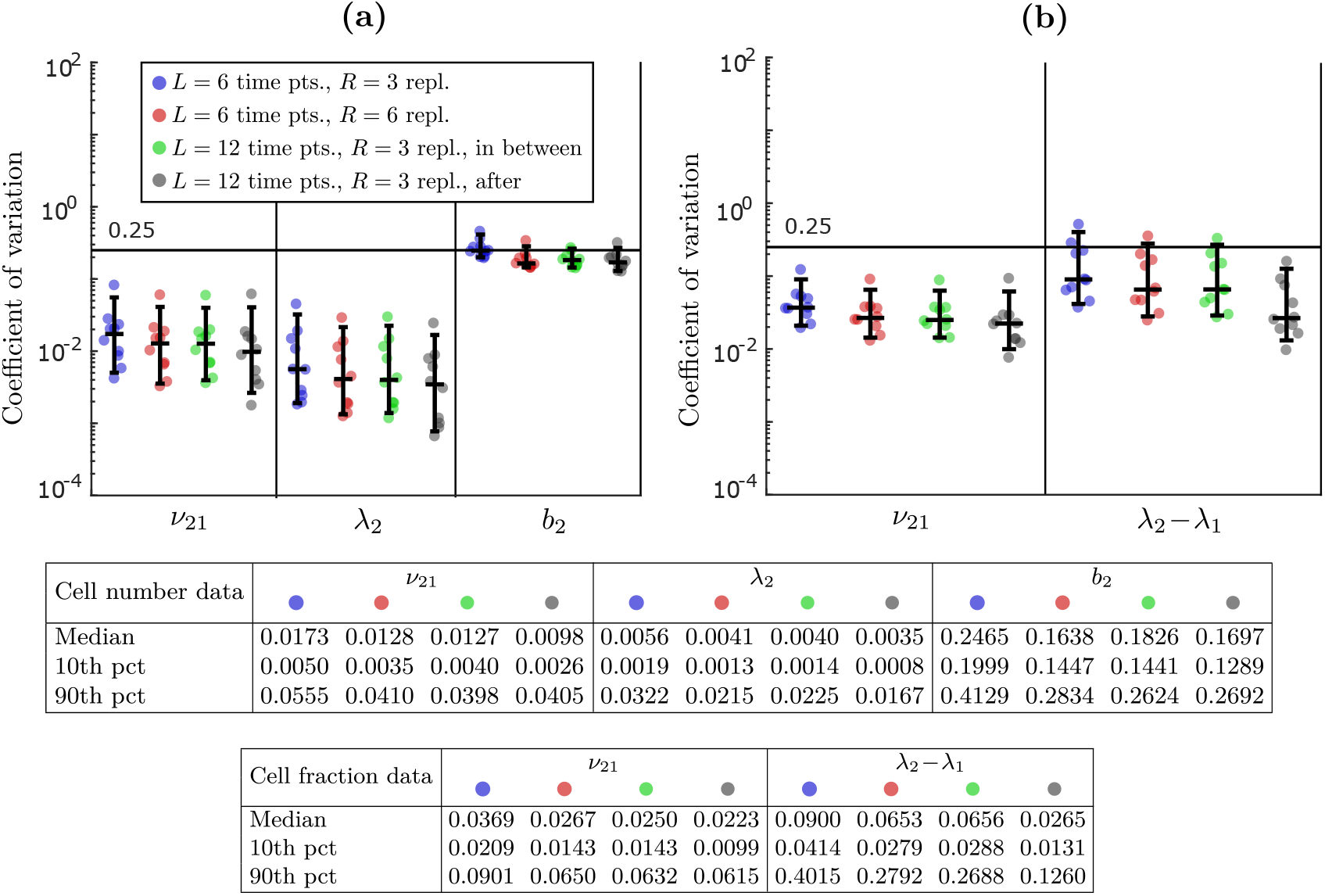
Comparison of estimation error for different experimental designs when the number of data points is doubled. We generated 10 parameter regimes and 100 datasets for each regime. The blue dots represent estimation from datasets with *L* = 6 time points and *R* = 3 replicates. The red dots represent estimation from *L* = 6 time points and *R* = 6 replicates. The green and grey dots represent estimation from *L* = 12 time points and *R* = 3 replicates, where the extra time points are added in between and after the previous time points, respectively. Panel **(a)** shows estimation from cell number data and panel **(b)** shows estimation from cell fraction data.

For cell number data, the median CV for the switching rate *ν*_21_ and the net birth rate *λ*_2_ reduces by 26% and 27%, respectively, when the number of replicates is doubled (design 1) (Fig. 5a). This is consistent with the fact that the standard deviation of an MLE estimator can be expected to decrease with 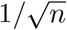, where *n* is the number of datapoints 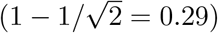 [42]. Adding data from time points in between the previous time points (design 2) has a similar effect on the median CV. However, adding time points after the previous points (design 3) reduces the median CV of *ν*_21_ and *λ*_2_ by 23% and 16%, respectively, over adding replicates (design 1). We also note that the 10th percentile of the CV for *ν*_21_ and *λ*_2_ reduces by 26% and 42%, respectively, between design 1 and design 3, which indicates that the degree of improvement between design 1 and design 3 depends very much on the parameter regime.

For cell fraction data, the relative attractiveness of the three experimental designs is similar (Fig. 5b). However, in this case, the estimate for the net birth rate difference *λ*_2_ − *λ*_1_ benefits significantly more from using design 3 than the estimate for the switching rate *ν*_21_.

For example, the median CV for *ν*_21_ reduces by 16% and the 10th percentile by 30% between design 1 and design 3, while the analogous reduction for *λ*_2_ − *λ*_1_ is 59% and 53%, respectively. In our structural identifiability analysis for cell fraction data (Section 6.2), we observed that it is more difficult to estimate *λ*_2_ − *λ*_1_ than *ν*_21_ from the initial population dynamics, and that *λ*_2_ − *λ*_1_ can be identified from the equilibrium proportions 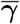 if the switching rates ***ν*** are known. The fact that adding more information on the long-run behavior of the population benefits the estimation of *λ*_2_ − *λ*_1_ more than *ν*_21_ is consistent with these insights. Of course, the results of the previous Section 7.3 indicate that the estimation of *λ*_2_ − *λ*_1_ can be improved even further by using cell number data as opposed to cell fraction data.

In general, Sections 7.3 and 7.4 show how our framework can be used to evaluate the estimation accuracy that can be achieved by different experimental designs, depending e.g. on what data is collected, when it is collected, how many replicates are performed, etc.

### 7.5 Improving identifiability of the rates of cell division and cell death

For cell number data, even though the birth rates **b** can be estimated reasonably well in many cases by Section 7.3, they are estimated much less accurately than the net birth rates ***λ*** and the switching rates ***ν***. In Figure 7a, we show that as the number of replicates is increased from 3 to 20 or above, the accuracy in the estimation becomes more acceptable. However, even with 100 replicates, the birth rates **b** are estimated less accurately than the net birth rates ***λ*** with 3 replicates (see Figure 4).

As we mentioned in the introduction, data on the number of cells in each state at each time point can be obtained by measuring the fraction of cells in each state and the total number of cells at each time point. In addition, it is often possible to measure the number of dead cells at each time point, see e.g. [30]. If these data are obtained, we can augment our mathematical model by introducing a new cell state, which cells transition into upon death (Figure 6). In Figure 7b, we show that if we apply our estimation framework to this model, the birth rates **b** become as easy to estimate as the net birth rates ***λ***. Thus, if data is collected on the number of live and dead cells at each time point, it becomes possible to estimate all model parameters accurately using our framework.

**Figure 6:**
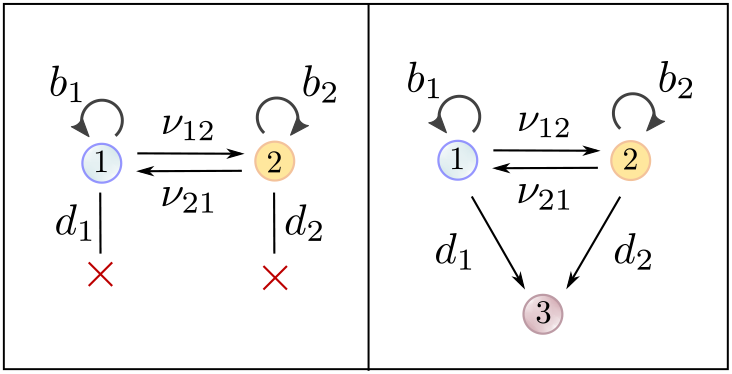
Augmentation of the mathematical model for when data is available on the number of dead cells at each time point. In that case, in stead of cells being lost from the model upon dying (left panel), they transition into a new state (right panel).

**Figure 7:**
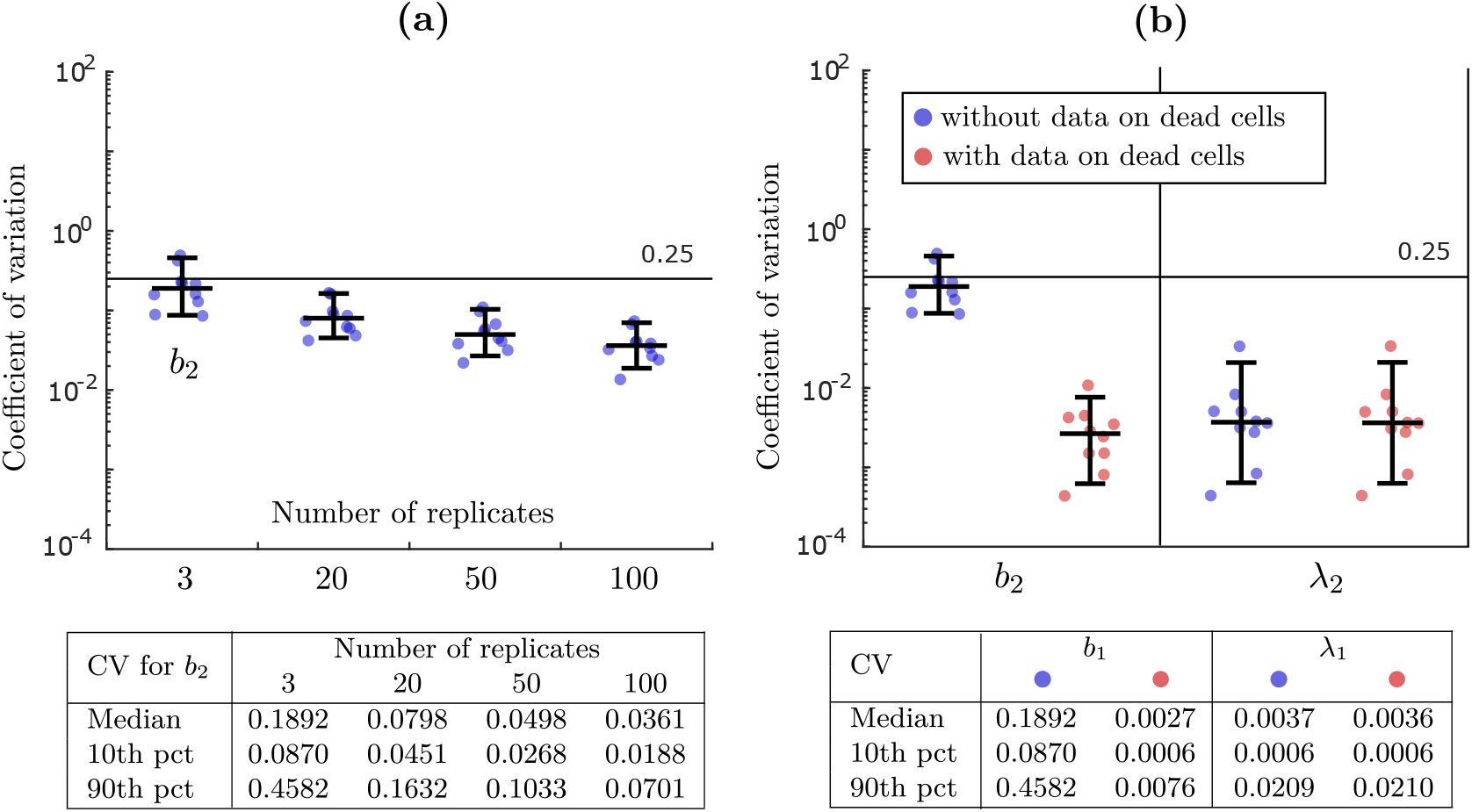
Two ways of improving the estimation accuracy for the birth rates **b** when cell number data is used. In **(a)**, we show how the estimation accuracy for the birth rate *b*_2_ improves as the number of experimental replicates is increased. In **(b)**, we compare the estimation accuracy for the birth rate *b*_2_ and the net birth rate *λ*_2_ depending on whether data on the number of dead cells at each time point is included in the estimation or not.

It should be noted that data collection on the number of dead cells is confounded by the fact that dead cells are eventually cleared from the system. This can potentially be addressed by introducing a clearance rate for dead cells in the augmented model, i.e. by introducing a death rate for the type-3 cells in Figure 6.

### 7.6 Estimation using endpoint data vs. sequential data

We conclude by examining how well our estimation framework applies to sequential data, when data is collected at multiple time points in the same experiment (Section 3). In Figure 8, we see that for cell number data, the CV for each parameter approximately doubles when applying our framework to sequential data vs. endpoint data. However, it remains true that the switching rates ***ν*** and net birth rates ***λ*** can be estimated with good accuracy. For cell fraction data, the difference in the estimation error for ***ν*** and *λ*_2_−*λ*_1_ is even smaller. Together, these results indicate that our framework can yield reasonable estimates for sequential data.

**Figure 8:**
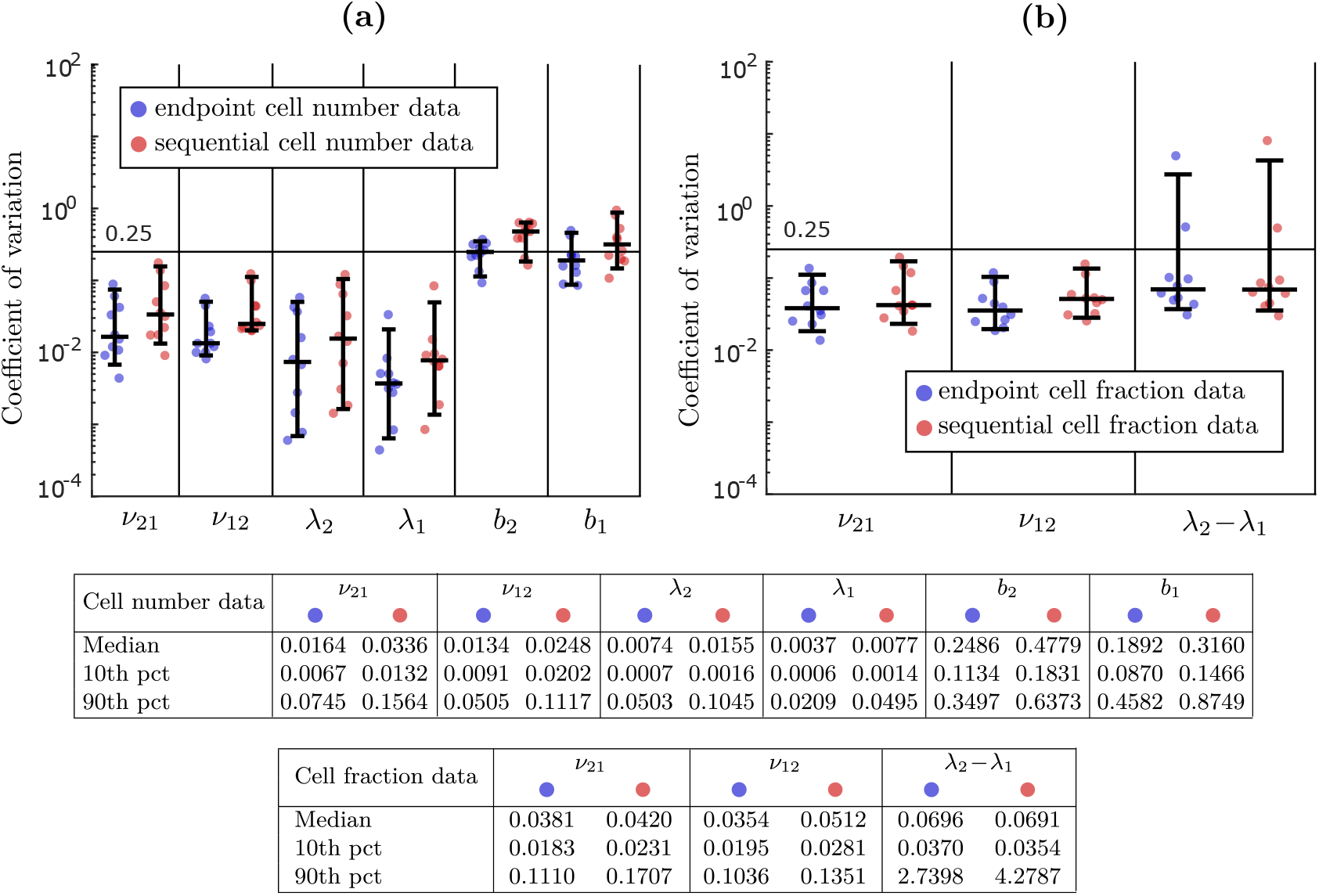
Comparison of estimation error depending on whether our framework is applied to endpoint data or sequential data. The blue dots show the estimation error when endpoint data is used, i.e. when experiments from different time points are independent, and the red dots show the error when sequential data is used, i.e. when data is collected at multiple time points in the same experiment. Panel **(a)** shows the comparison for cell number data and panel **(b)** for cell fraction data. Even though our framework is derived for endpoint data, it provides reasonable estimation accuracy for sequential data.

At the same time, for cell number data in particular, there can be a significant benefit to developing a method tailored to sequential data.

## 8 Application: Transition between stem and non-stem cell states in SW620 colon cancer

To give an example of how our estimation framework can be used to analyze experimental data, we conclude by applying it to a publicly available cell fraction dataset. We use data collected by Yang et al. [17] and made available in Tables S2 and S3 of Wang et al. [20], on the dynamics between stem-like (type-1) and non-stem (type-2) cells in SW620 colon cancer. In Yang et al. [17], the two cell types were sorted based on expression of the CD133 cell-surface antigen marker. Isolated subpopulations were expanded and phenotypic proportions were tracked for 24 days, with data collected every other day. This dataset has previously been analyzed using the CellTrans estimation method [26] (Appendix A).

Since data on individual experimental replicates are not available, we use data on the mean cell fraction across replicates as input to our estimation framework. We first consider the statistical model (11) with 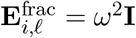 for all *i, l*, which we refer to as Model I:

### Model I: 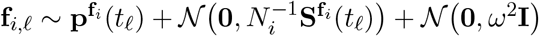

In Figure 9, we show parameter estimates and 95% confidence intervals (CIs) for *ν*_21_, *ν*_12_ and *λ*_2_ − *λ*_1_ under Model I. The CIs show that while the point estimates for *ν*_21_ and *ν*_12_ are 0.1540 and 0.0570, the true value of *ν*_21_ may range between 0.1110 and 0.2119, and the true value of *ν*_12_ may range between 0.0361 and 0.0872. Since the two CIs do not overlap, we can be confident that *ν*_21_ *> ν*_12_, but there is significant uncertainty as to the true values of these parameters. The CI for *λ*_2_ − *λ*_1_ is even wider, which is in line with our earlier observations that this parameter is more difficult to estimate from cell fraction data than the switching rates, especially when *λ*_2_ − *λ*_1_ is relatively small in magnitude (Sections 6.2 and 7.3). In fact, the CI for *λ*_2_ − *λ*_1_ includes zero, meaning that it is plausible that *λ*_1_ = *λ*_2_.

**Figure 9:**
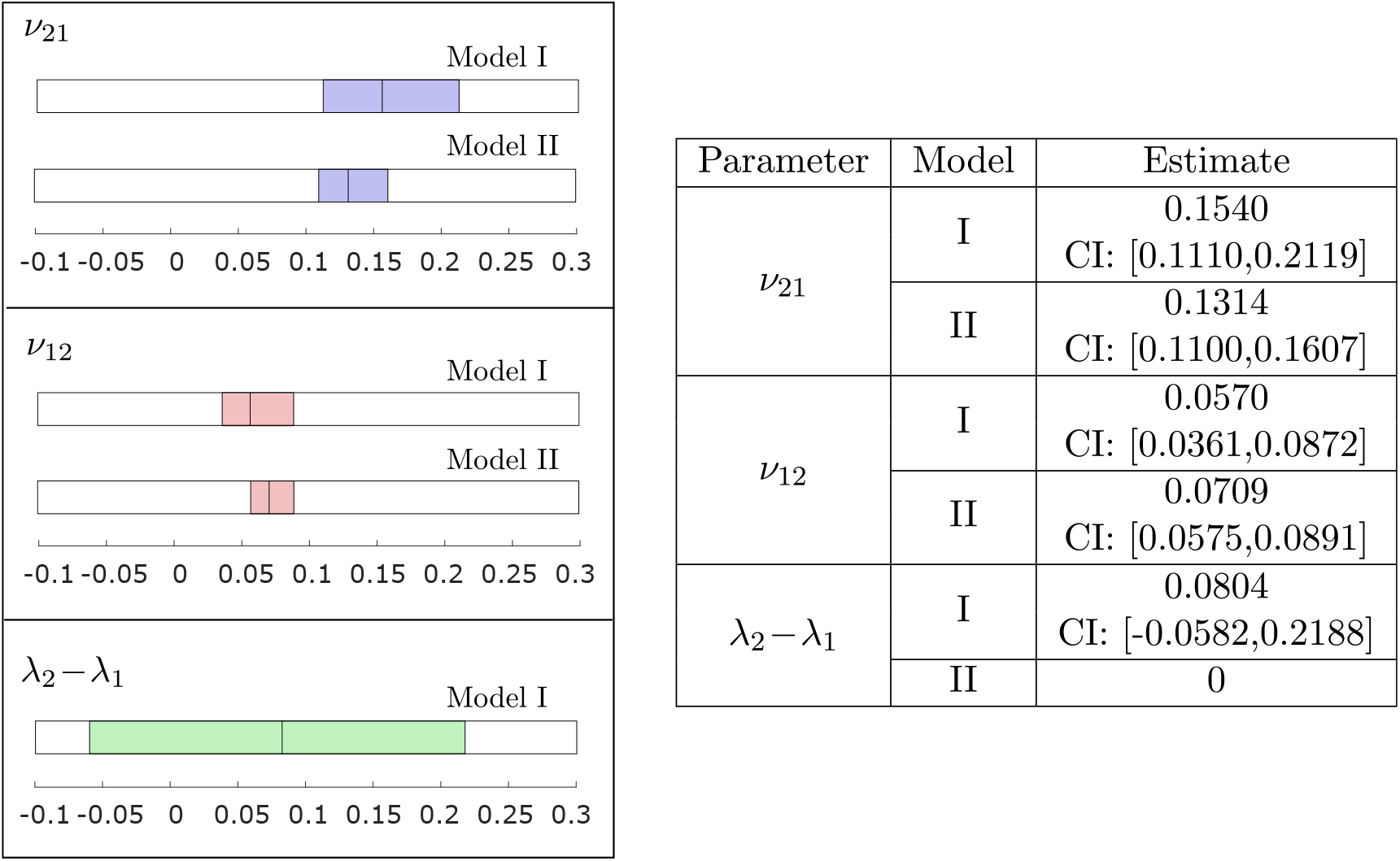
Comparison of point estimates and 95% confidence intervals for the statistical model 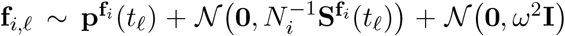 (Model I) and the same model with *λ*_2_ − *λ*_1_ = 0 (Model II) applied to publicly available cell fraction data from Yang et al. [17].

In the CellTrans paper [26], it is assumed that the two phenotypes have the same growth rates, based on measurements from Wang et al. [20]. We can build this assumption into our estimation by solving the MLE problem (14) for Model I under the constraint *λ*_2_ − *λ*_1_ = 0 (Appendix C). We refer to this as Model II:

### Model II: 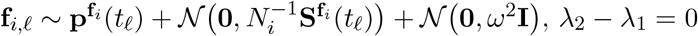

Estimation results for Model II are shown in Figure 9. The assumption *λ*_1_ = *λ*_2_ has a noticeable effect on both the point estimates of *ν*_21_ and *ν*_12_ and their confidence intervals. For example, the ratio *ν*_21_*/ν*_12_ is 2.70 under Model I, while it is 1.85 under Model II. In other words, switching from type-2 to type-1 happens about three times as often as switching from type-1 to type-2 under Model I, while it happens about two times as often under Model II. Furthermore, under Model II, the length of the CI for *ν*_21_ is reduced by a half compared to Model I, meaning that Model II significantly restricts the plausible values of *ν*_21_.

In the CellTrans paper [26], the same dataset is used to estimate switching probabilities of *p*_21_ = 0.1030 and *p*_12_ = 0.0545, based on a discrete-time Markov model with a time step of Δ*t* = one day. We also solved the TRANSCOMPP problem (16) (Appendix A) with Δ*t* = one day to obtain the estimates *p*_21_ = 0.1360 and *p*_12_ = 0.0537 for the switching probabilities and Λ_22_*/*Λ_11_ = 1.0787 for the ratio between the growth factors of the two phenotypes, which translates to a growth rate difference of *r*_2_ −*r*_1_ = 0.0758 if we set 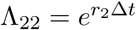 and 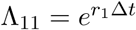. In the CellTrans and TRANSCOMPP models, type switches are synchronized between all cells in the population, and they occur at discrete time steps. In our continuous-time model, the time steps are infinitesimally small, and each cell has a certain probability of switching, proliferating and dying during each step, independently of other cells (Section 2.1). If we shorten the time step to Δ*t* = 1*/*10 day, the switching probabilities become 0.01111 and 0.00588 under CellTrans, which translates to continuous-time rates of 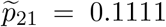 and 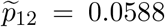. These estimates fall at the lower limits of our CIs for *ν*_21_ and *ν*_12_ under Model II (Figure 9). Under TRANSCOMPP, the switching probabilities become 0.01544 and 0.00570 for Δ*t* = 1*/*10 day, which translates to continuous-time rates of 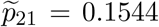 and 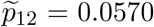, and the difference in growth rates becomes 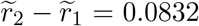. These estimates are very similar to the point estimates of Model I (Figure 9).

The estimates of CellTrans and TRANSCOMPP are consistent with our estimates in that they fall within the 95% confidence intervals produced by our framework, when the time step is taken to be sufficiently small. Our framework complements these methods for cell fraction data by enabling more complete analysis of the estimates and the uncertainty involved. First of all, estimates from discrete-time models are sensitive to the length of the time step chosen, which makes their interpretation less clear. Second, our framework provides likelihood-based confidence intervals, which encompass all of the plausible values for the model parameters. For example, the CIs for Model I reveal how uncertain the value of *λ*_2_ − *λ*_1_ is compared to *ν*_21_ and *ν*_12_. If only point estimates were reported for Model I, it would not be clear that *λ*_2_ − *λ*_1_ cannot be distinguished from zero, and that *λ*_2_ − *λ*_1_ may in fact be negative even though the point estimate is positive. Furthermore, the CIs for Model II show that even under the assumption *λ*_1_ = *λ*_2_, the true values of *ν*_21_ and *ν*_12_ may be around 50% larger than what is predicted by CellTrans. Third, under our framework, it is simple to incorporate assumptions such as *λ*_1_ = *λ*_2_ or *ν*_12_ = *ν*_21_ into the estimation (Appendix C) and to assess their effect on point estimates and CIs. As we have observed, the assumption *λ*_1_ = *λ*_2_ significantly restricts the plausible values of *ν*_21_ and *ν*_12_, which may underestimate the true uncertainty in the estimation, given that the claim *λ*_1_ = *λ*_2_ is subject to statistical error. Finally, of course, if data is collected on the total size of the population at each time point, our framework can make use of the added data to produce improved parameter estimates.

## 9 Discussion

In this work, we have proposed a maximum likelihood framework for estimating the rates of cell proliferation and phenotypic switching in cancer. In contrast to previous approaches, the framework explicitly models the stochastic dynamics of cell division, cell death and phenotypic switching, it provides likelihood-based confidence intervals for the model parameters, and it enables estimation from data on the fraction of cells or the number of cells in each state at each time point. An implementation of our framework in MATLAB with sample scripts is available at https://github.com/egunnars/phenotypic_switching_inference/.

We have also used our framework to analyze the identifiability of model parameters. Through a combination of theoretical and numerical investigation and application to real data, we have seen that when cell fraction data is used, the switching rates ***ν*** may be the only parameters that can be estimated accurately, while the net birth rate differences ***λ***^[−1]^ may also be obtainable when they are sufficiently large. Including information on the total size of the population at each time point yields significantly better estimates of ***λ***^[−1]^, and it also enables accurate estimation of the net birth rates ***λ***. Finally, if enough experimental replicates are performed, or if data is collected on the number of dead cells at each time point, it even becomes possible to estimates the birth rates **b** and death rates **d** accurately. In a previous work, we discussed how knowledge of the model parameters ***ν, λ*, b** can enhance our understanding of resistance evolution in cancer and inform the design of combination treatments of anti-cancer agents and epigenetic drugs [25]. Together, these parameters shape the evolution of phenotypic proportions and the tumor burden over time, each of which is relevant to the dynamics of tumor recurrence. Our current work shows that it is not possible to estimate the net birth rates ***λ*** or the birth rates **b** accurately from cell fraction data, it indicates what data is required to obtain these parameters, and it offers a rigorous approach to parameter estimation and uncertainty quantification once the data has been acquired.

There are several avenues for future development of the framework. First, our multitype branching process model assumes that cells are allowed to grow uninterrupted for the duration of the experiments. This does not address the effect of passaging in longer-duration experiments. One possible solution is to keep track of cell state proportions and seeding densities for each passage, and treat each new passage as a new experiment. However, our framework currently assumes that initial conditions are known, while uncertainty is assigned to all subsequent time points. In reality, the initial conditions are subject to measurement error, and it may become important to model this error for the case of repeated passaging.

Second, our framework currently models measurement error as an additive Gaussian noise with a general covariance matrix. We have suggested simple ways of choosing the covariance matrix both for cell number and cell fraction data, but further exploration of appropriate choices is warranted. Depending on the application, it may also be necessary to develop a more sophisticated model for the measurement error. For example, for cell number data, if the measurement error is proportional to the population size, it may be necessary to model it as a multiplicative term rather than an additive term, or to build the experimental cell counting procedure more explicitly into the statistical model. We plan to address this in future work.

Third, we have focused on estimation from experiments started with isolated subpopulations of each phenotype, as this is a common experimental design, and we have analyzed parameter identifiability in this setting. Understanding to what extent the model parameters, or some combinations of the parameters, can be estimated from more limited data is an interesting avenue for future investigation. For example, if we are interested in estimating parameters from clinical data, the data will likely contain much less information than we have assumed here, and it will become necessary to analyze what parameters are identifiable and how identifiability can be improved, e.g. by combining data from similar patients.

Finally, we believe our framework can be useful for the design of cell line experiments aimed at deciphering the dynamics of phenotypic switching. For example, preliminary experiments can first be conducted, from which initial parameter estimates and confidence intervals can be derived. Based on the confidence intervals, one can construct a set of likely values for the parameters, which can be used to evaluate the expected improvement in estimation accuracy depending on the experimental design (see e.g. [43]). Once good experimental designs have been identified, one can evaluate whether the expected improvement in estimation accuracy justifies the additional experimental resources. If this is the case, additional experiments can be performed and the process can be repeated. In a future work, we plan to develop a tool for the optimal selection of experimental designs, to facilitate more efficient utilization of experimental resources.

## A Review of existing methods

At the single-cell-level, phenotypic switching has commonly been modeled by a discrete-time Markov chain with *K* ≥ 2 states, where *K* is the number of phenotypes. In each time step, a cell in state *j* transitions to state *k* ≠ *j* with probability *p*_*jk*_, and it remains in state *j* with probability *p*_*jj*_ = 1 − Σ_*k*≠*j*_*p*_*jk*_. The transition probabilities are collected into the *K × K transition matrix* **P** = (*p*_*jk*_). The evolution of the Markov chain is determined by **P** and the *initial distribution* **q** = (*q*_1_, …, *q*_*K*_), where *q*_*j*_ is the probability that a cell starts in state *j*. If we let **q**^(*ℓ*)^ denote the cell state distribution after *ℓ* ≥ 1 time steps, then **q**^(*ℓ*)^ = **qP**^*ℓ*^.

Say we conduct *K* cell line experiments starting with *N* cells in each experiment and known initial cell state distributions **q**_1_, …, **q**_*K*_. The initial distributions are collected into a *K × K* matrix **Q**, where **q**_*i*_ is the *i*-th row vector. Each experiment is run for *ℓ* ≥ 1 time steps, at which point the fraction of cells in each state is recorded. Let 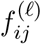 be the observed fraction of cells in state *j* under the *i*-th initial condition. The observations at the *ℓ*-th time step under the *i*-th initial condition are collected into a vector 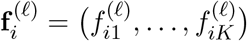, and all observations at the *ℓ*-th time step are collected into a *K × K* matrix 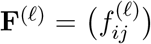. If there are multiple replicates *r* = 1, …, *R*, we let **F**^(*ℓ*),*r*^ denote the data from the *r*-th replicate.

Now, assume that the starting population *N* is large, that there is no cell division or cell death, and that each cell switches between states according to the above Markov model. In this case, by the law of large numbers, the model-predicted distribution between cell states **QP** after *ℓ* time steps can be approximated by the experimentally observed cellstate fractions **F**^(*ℓ*)^. If we equate these two matrices, we can obtain an estimate **P** of the transition matrix **P** by inverting the matrix **Q** of initial distributions and taking an *ℓ*-th matrix root, **P** = (**Q**^−1^**F**^(*ℓ*)^) ^1/*ℓ*^. Here, we assume that **Q** is invertible, which is e.g. the case when experiments are started with isolated subpopulations.

This simple estimation idea was applied by Gupta et al. [21] to investigate phenotypic switching between stem-like, basal and luminal cell states in breast cancer, using data from a single time point. A multiple-time-point extension has since been implemented in the R package CellTrans [26]. Say that cell state fractions are experimentally observed at time steps *m*_1_, …, *m*_*L*_ for *L* ≥ 1. CellTrans first computes an estimate 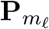 of the transition matrix for each time step as above, and then returns a final estimate as the average across time steps:

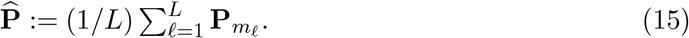

CellTrans also involves a regularization step to ensure that 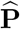 is stochastic. CellTrans is used on publicly available datasets in [26] and it has been applied more recently in [44, 32, 45].

Cell populations in culture typically change in size over time. The above idea can be applied to growing populations assuming that all phenotypes grow at the same rate and that a constant-sized model is thus sufficient to describe the evolution of cell state fractions. Both Gupta et al. [21] and Su et al. [12] have applied an augmented version of the above model which is intended to capture proliferation differences between types. In the augmented model, during a single time step, each type-*j* cell first grows deterministically to a population of size Λ_*jj*_, and a fraction *p*_*jk*_ of cells then switch to type-*k*. The growth factors Λ_*jj*_ are collected into a diagonal proliferation matrix **Λ**, and the multiple **ΛP**, normalized to produce cell fractions as opposed to cell numbers, is used to predict the distribution between cell states. In both Gupta et al. [21] and Su et al. [12], the matrix **Λ** is found by randomly sampling candidate parameter values and selecting the values that best fit the experimental data.

TRANSCOMPP [27] is a more systematic version of the above method. In TRANSCOMPP, the diagonal proliferation matrix **Λ** and the transition matrix **P** are estimated by minimizing the sum of squared errors between the model prediction and the data,

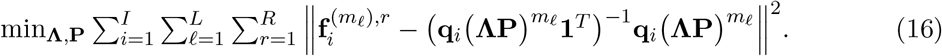

Note that only the relative growth factors Λ_*jj*_*/*Λ_11_ for *j* = 2, …, *K* are identifiable from this problem. TRANSCOMPP is implemented in MATLAB, and it includes a stochastic resampling procedure for estimating the distributions of the transition probability estimates.

In modeling switching between HER2+ and HER2− states in breast cancer, Li and Thirumalai [28] employ a deterministic continuous-time model. Their model assumes symmetric and asymmetric cell divisions, which through reparametrization leads to the same dynamics as symmetric cell divisions and switching between types. Li and Thirumalai assume equal rates of asymmetric division for the two types (or equivalently, equal rates of switching between types), and they show that if experiments are started with isolated subpopulations, the slopes of the cell fraction trajectories at time 0 can be used to estimate these rates. They also show that the equilibrium proportion between types can be used to estimate the difference in symmetric division rate between the two types. The proportion between phenotypes in the parental population is used as an estimate of the equilibrium proportion. We have made use of these insights in our identifiability analysis in Section 6.2 of the main text.

Finally, in their investigation of epithelial to mesenchymal transition in breast cancer, Devaraj and Bose [30, 46] employ a discrete-time model where cells divide, die and switch between types. Their model includes a separate state for dead cells to facilitate estimation of death rates and well as division rates. We have used the same idea in Section 7.5 to improve the identifiability of birth and death rates under our framework. Their model furthermore assumes that the rates of birth, death and switching are time-dependent. Devaraj and Bose derive difference equations for the change in the number of cells in each state between time points. They then propose a multi-objective optimization problem to estimate the model parameters from data on phenotypic proportions and the number of alive and dead cells at each time point. Their parameter fitting procedure minimizes the least squares error between the model predictions and the data across the different time points, while ensuring that parameters do not vary too drastically between time periods.

## B Estimation for reducible switching dynamics

In the main text, we have assumed that the switching dynamics are irreducible, meaning that it is possible to switch between any pair of phenotypes, possibly through intermediate types. In this section, we show how our framework can be applied to the case of reducible switching dynamics. For simplicity, we will consider one particular model shown in Figure 10. This model has been applied e.g. to the dynamics of epigenetic gene silencing under recruitment of chromatin regulators [47] and the evolution of epigenetically-driven drug resistance in cancer, where drug-sensitive cells (type-1) first acquire a transiently resistant phenotype (type-2) and then evolve to stable epigenetic resistance (type-3) [25].

**Figure 10:**
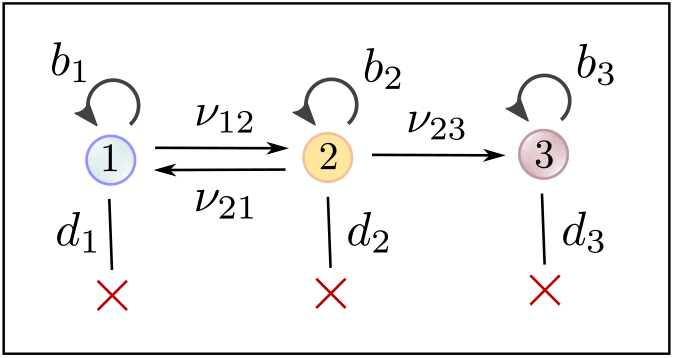
To demonstrate that our estimation framework is applicable to reducible switching models, we consider a three-type model with a reversible transition between type-1 and type-2, and an irreversible transition from type-2 to type-3. This model is applicable e.g. to epigenetic gene silencing under the recruitment of chromatin regulators [47] and to epigenetically-driven drug resistance in cancer [25].

Say that experiments are conducted from isolated initial conditions, and say first that cell number data is collected. For the model in Figure 10, the distribution of the data vector **n**_3,, *r*_ is degenerate, since *n*_3,, *r,j*_ = 0 for *j* = 1, 2. As a result, the covariance matrix **Σ**^(3)^(*tℓ*) is singular for all *ℓ* = 1, …, *L*, and the likelihood function in (6) is not defined. To resolve this issue, we set **C**_1_ = **C**_2_ = **I** and 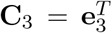, where **e**_3_ is the 1 *× K* third unit vector. By Proposition 1, **n**_3,*ℓ,r*_**C**_3_ = *n*_3,*ℓ,r*,3_ has a normal distribution, which is nondegenerate. We therefore modify the likelihood function in (6) to

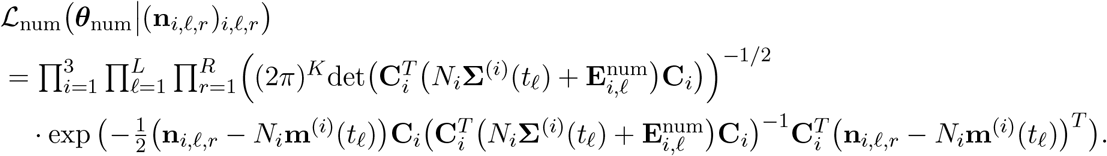

From this likelihood function, MLE estimates and confidence intervals can be computed as described in Section 4.3, where we restrict the set of feasible parameters **Θ**_num_ so that *ν*_13_ = *ν*_31_ = *ν*_32_ = 0. By our analysis in Section 6.1, all model parameters are structurally identifiable for this example.

To accommodate model structures such as the one discussed here, the above modified likelihood function is implemented in our MATLAB codes (Appendix C). By taking **C**_*i*_ = **I** for each *i* = 1, …, *I*, we recover the original likelihood function in (6).

If cell fraction data is collected, there is no value in conducting experiments starting only from type-3 cells. We therefore use the likelihood function

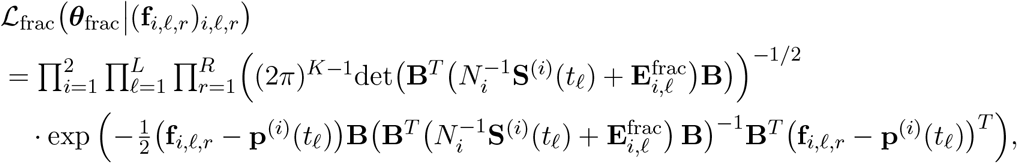

where we only include experiments started by type-1 and type-2 cells, respectively. By our analysis in Section 6.2, the switching rates *ν*_12_, *ν*_21_ and *ν*_23_, and the net birth rate differences *λ*_2_ −*λ*_1_ and *λ*_3_ −*λ*_2_, are structurally identifiable in this case. An example of a model structure where it becomes necessary to modify the above likelihood function for cell fraction data is given in Appendix C.

## C Implementation in MATLAB

In this section, we discuss how our estimation framework is implemented in MATLAB.

### C.1 Cell number data

The first step in the implementation for cell number data is to compute simple parameter estimates for the switching rates ***ν*** and the net birth rates ***λ*** based on a deterministic population model. This model is obtained by ignoring the stochastic terms in the statistical model (5), i.e. by equating the data vector **n**_*i,ℓ,r*_ with the mean prediction of (5):

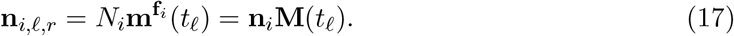

Let **N** be the *I × K* matrix with the initial conditions **n**_*i*_ as row vectors, and let **N**_*ℓ,r*_ be the *I × K* matrix with the data vectors **n**_*i,ℓ,r*_ as row vectors. We can then write (17) in matrix form as

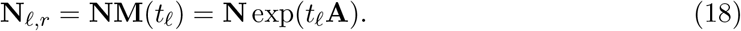

Assuming **N** has rank *K*, we can solve for **A** in (18) by first multiplying both sides by **N**^*T*^, then multiplying both sides by the inverse of **N**^*T*^ **N**, and finally taking a matrix logarithm. We can thus obtain an estimate for the infinitesimal generator **A**,

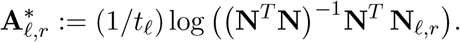

We then compute a final estimate **A**^*\**^ by averaging across time points and replicates:

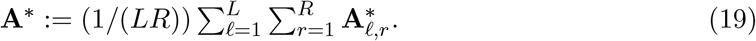

From **A**^*\**^, we can obtain estimates of the switching rates ***ν*** and the net birth rates ***λ***.

As indicated in Appendix B, we implement the following likelihood function in our codes:

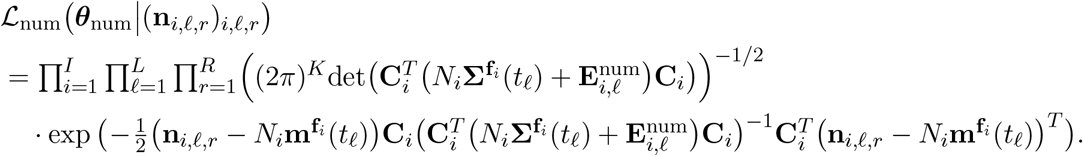

For each *i* = 1, …, *I*, **C**_*i*_ is a *K × J*_*i*_ matrix for some 1 ≤ *J*_*i*_ ≤ *K*, which can be used to reduce the dimension of the data vector **n**_*i,ℓ,r*_ when necessary. This option can e.g. be useful for models with reducible switching dynamics, see Appendix B.

From the above likelihood function, we compute a negative double log-likelihood as in (7), and solve the MLE problem (8) using the sequential quadratic programming (sqp) solver in MATLAB. For the optimization, one must supply an initial guess 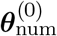 for the parameter vector ***θ***_num_, and a set of feasible parameters **Θ**_num_ of the form

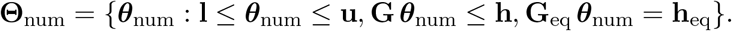

By default, we assume lower bounds of **0** for the switching rates ***ν*** and the birth rates **b**, and we impose the inequality constraint ***λ*** ≤ **b**. The user is expected to provide lower bounds for the net birth rates ***λ*** and upper bounds for all parameters, and they have the option to provide further inequality or equality constraints as necessary. This provides the opportunity to impose constraints such as *λ*_1_ = *λ*_2_ (Section 8) or *ν*_13_ = *ν*_31_ = *ν*_32_ = 0 (Appendix B).

For the initial guess 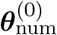, we use the simple estimates for ***ν*** and ***λ*** computed from (19). An initial guess for the birth rate *b*_*i*_ is generated as |*λ*_*i*_|*/U*, where *U* is uniformly distributed between 0 and 1. The idea is that if *λ*_*i*_ *>* 0, then in the absence of phenotypic switching, the survival probability of a single-cell derived clone of type *i* is *q*_*i*_ = *λ*_*i*_*/b*_*i*_ [48]. Since we do not assume any information on *q*_*i*_, we sample it uniformly between 0 and 1, and then use the initial guess for *λ*_*i*_ to compute an initial guess for *b*_*i*_.

If data on the number of dead cells at each time point is available, the initial guesses for the birth rates can be improved as follows. As before, let *n*_*ij*_ be the number of starting cells of type-*j* under the *i*-th initial condition. In the absence of phenotypic switching, the expected number of type-*j* cells at time *t* under the *i*-th initial condition is given by *n*_*ij*_ exp(*λ*_*j*_*t*). If we assume that type-*j* cells grow deterministically according to this function, the number of dead cells of type-*j* that accumulate up until the first experimental timepoint *t*_1_ is given by

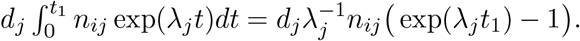

Set 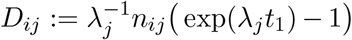 and let **D** = (*D*_*ij*_) denote the corresponding *I × K* matrix. Also, let **c** denote the 1 *× I* vector of the experimentally measured number of dead cells at time *t*_1_, averaged across the *R* experimental replicates. We should then have

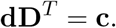

Assuming **D** has rank *K*, we can solve this equation for **d** as follows:

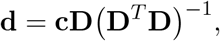

which gives an estimate for the vector of death rates **d**. An estimate for the birth rates **b** can then be computed as **b** = ***λ*** + **d**.

In addition to being used to initialize the optimization, the initial guess 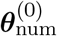 is used to estimate the relative scales of the parameters ***ν, λ*** and **b**. In particular, for the *i*-th coordinate of the initial guess, we define the corresponding scale variable

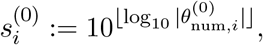

with 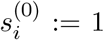 if 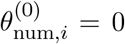. For example, if the initial guesses are **b**^(0)^ = (1.5, 1.2) for the birth rates, ***λ***^(0)^ = (0.3, 0.4) for the net birth rates, and ***ν***^(0)^ = (0.05, 0.002) for the switching rates, the corresponding scale variables are (1, 1), (0.1, 0.1) and (0.01, 0.001), respectively. For a given parameter vector ***θ***_num_, we define the transformed vector

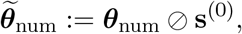

where denotes elementwise division. For the initial guesses **b**^(0)^ = (1.5, 1.2), ***λ***^(0)^ = (0.3, 0.4) and ***ν***^(0)^ = (0.05, 0.002), the corresponding transformed values are 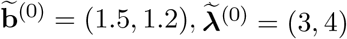 and 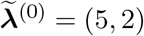. With this transformation, all nonzero parameters take values in [1, 10]. When we solve the MLE problem (8), we treat 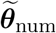 as the parameter vector instead of ***θ***_num_, and solve

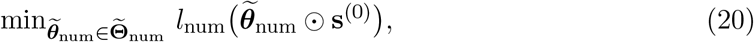

where 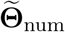 is the transformed set of feasible parameters. The parameter scaling is applied to ensure that all model parameters are of a similar magnitude in the optimization.

In most cases, we have found it sufficient to solve the optimization problem (20) once. However, in our codes, we provide an option to solve the problem multiple times, using (i) user-supplied initial guesses, (ii) initial guesses based on the simple estimates from (19), with new birth rates selected randomly each time, or (iii) randomly sampled initial guesses, using the parameter generation procedure described in Appendix D below.

The optimization problems (9) for the endpoints of the confidence intervals are solved in a similar way, except the initial guess is taken to be the maximum likelihood estimate.

### C.2 Cell fraction data

The implementation for cell fraction data is similar with the following modifications. First of all, we parametrize the model in terms of the death rates **d**, the net birth rate *λ*_1_ and the net birth rate differences ***λ***^[−1]^, instead of the birth rates **b** and net birth rates ***λ*** (Section 5.2). Second, the initial guess for the MLE problem (14) is based on solving the following least squares problem, which minimizes the sum of squared errors between the mean prediction of the statistical model (11) and the data:

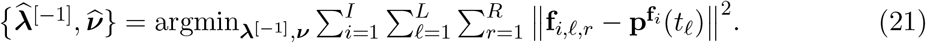

Note that this is a continuous-time version of the TRANSCOMPP problem (16). When solving (21), we need to supply an initial guess. If experiments are conducted from isolated initial conditions, we compute initial guesses for the switching rates ***ν*** based on part (1) of Proposition 4, which shows how ***ν*** can be estimated from the slopes of the mean functions **p**^(*j*)^(*t*) at time zero. We approximate the slopes of **p**^(*j*)^(*t*) at time zero using experimentally observed cell fractions at the first time point. The initial guesses for the remaining parameters are set to 0. If experiments are not conducted from isolated initial conditions, we randomly sample initial guesses as described in Appendix D below. The simple problem (21) returns estimates for ***ν*** and ***λ***^[−1]^, which we use as initial guesses for (14).

In our codes, we implement the following likelihood function for cell fraction data:

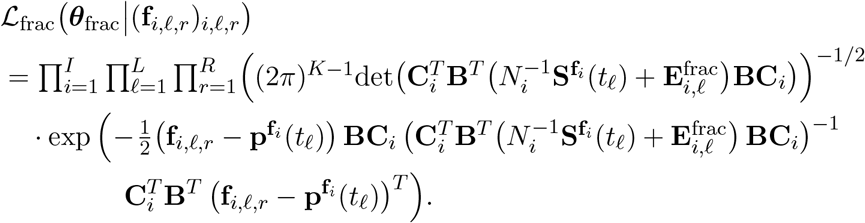

Recall from (12) that the matrix **B** is applied to reduce the data vector **f**_*i,ℓ,r*_ to a (*K* − 1)-dimensional vector. To accommodate reducible switching dynamics, the user is allowed to implement a further reduction in the data by specifying a (*K* − 1) *× J*_*i*_ matrix **C**_*i*_ for each initial condition *i*. This can for example be useful for the four-type model (*K* = 4) displayed in Figure 11, in which case we would take *I* = 3, **C**_1_ = **C**_2_ = **I** and 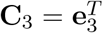, and we would restrict the set of feasible parameters **Θ**_frac_ so that *ν*_13_ = *ν*_14_ = *ν*_24_ = *ν*_31_ = *ν*_32_ = *ν*_41_ = *ν*_42_ = *ν*_43_ = 0. Note that here, **I** refers to the (*K* − 1) *×* (*K* − 1) = 3 *×* 3 identity matrix.

**Figure 11:**
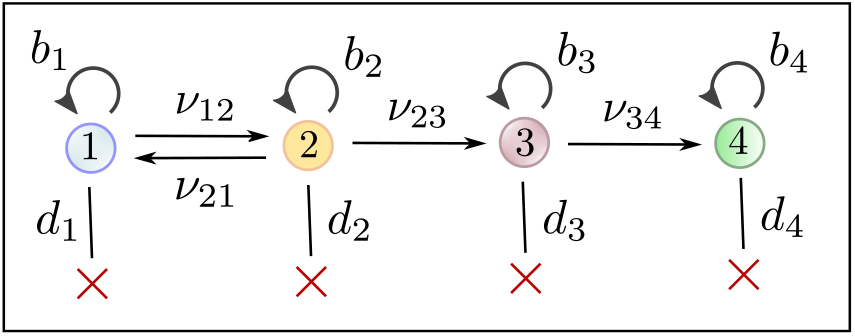
An example of a four-type switching model where the likelihood function (12) for cell fraction data from the main text must be modified to avoid degeneracy issues. This model structure can e.g. arise in the context of epigenetically-driven drug resistance in cancer, where drug-sensitive (type-0) cells can acquire transient resistance (type-1), which then evolves gradually to stable resistance (type-4) in two steps [25].

## D Generation of artifical data

Here, we discuss how the artificial data was generated for the numerical experiments in Section 7. First, to generate each parameter regime, we sampled the birth rates **b** and death rates **d** uniformly at random on (0, 1), with the following caveats: The birth rates **b** and net birth rates ***λ*** were required to be larger than 0.01 in absolute value, and at least one of the net birth rates *λ*_1_, …, *λ*_*K*_ was required to be positive. Each switching rate *ν*_*ij*_ was sampled as 10^−3+2*U*^, where *U* is uniform between 0 and 1, meaning that it was sampled log-uniformly between 10^−3^ and 10^−1^. The starting number of cells *N*_*i*_ was chosen as *N*_*i*_ = 10^−3^, *N*_*i*_ = 10^−4^ or *N*_*i*_ = 10^−5^ for *i* = 1, …, *K* based on the order of magnitude of the smallest switching rate. The experimental time points were selected as *t* = 1, …, 6.

In Section 7.4, where the number of time points was doubled, the time points were taken as either *t* = 0.5, 1, 1.5, 2, …, 6 or *t* = 1, 2, 3, …, 12, depending on whether the new time points were added in between or after the previous time points.

Once the parameters were set, we performed stochastic simulations of the model in Section 2 to obtain the artificial datasets. MATLAB codes for generating the parameter regimes and artifical datasets, and for performing estimation on the artifical datasets, are available at https://github.com/egunnars/phenotypic_switching_inference/.

## E Proof of Proposition 1

*Proof of Proposition 1*. First note that we can write

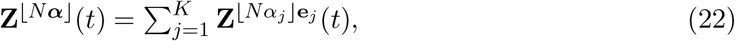

where 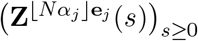 for *j* = 1, …, *K* are independent branching processes started with ⌊ *Nα*_*j*_ ⌋ cells of type-*j*, respectively. For each process, we can write

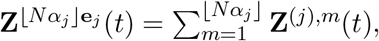

Where (**Z**^(*j*),*m*^(*s*))_*s*≥0_ for *m* = 1, …, ⌊ *Nα*_*j*_ ⌋ are i.i.d. copies of the branching process (**Z**^(*j*)^(*s*))_*s*≥0_ started by a single type-*j* cell. Set

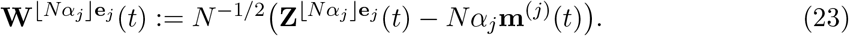

Let *J* ≥ 1 and let **C** be a *K × J* matrix. By the (multivariate) central limit theorem, as *N* → ∞,

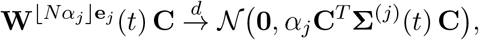

where **Σ**^(*j*)^(*t*) is the covariance matrix for **Z**^(*j*)^(*t*). We can then conclude from (22) that as *N* → ∞,

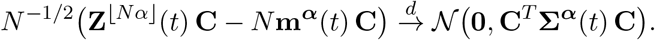

It remains to derive the expression (4) for the covariance matrix **Σ**^(*j*)^(*t*). To that end, let **D**^(*j*)^(*t*) be the matrix of second factorial moments of **Z**^(*j*)^(*t*),

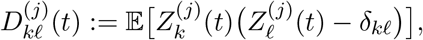

where *δ*_*kl*_ is the Kronecker delta. Let **s** = (*s*_1_, …, *s*_*K*_) be a *K*-dimensional vector of real numbers and set *h*_*j*_ := *b*_*j*_ + *d*_*j*_ + *k*≠*j ν*_*jk*_ for *j* = 1, …, *K*. Furthermore, let

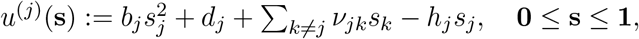

be the infinitesimal generating function for **Z**^(*j*)^(*t*), and let

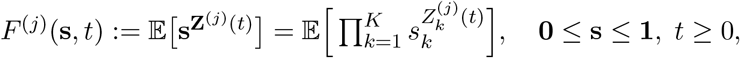

be the probability generating function for **Z**^(*j*)^(*t*). With this notation, we can write the Kolmogorov forward equation for **Z**^(*j*)^(*t*) as

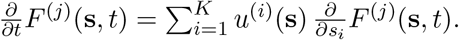

Then, for *k, ℓ* = 1, …, *K*,

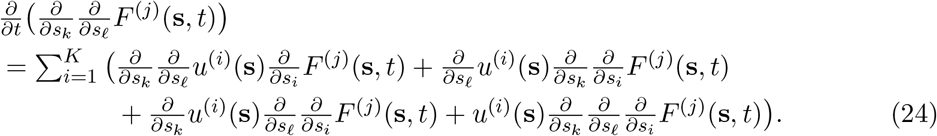

Now,

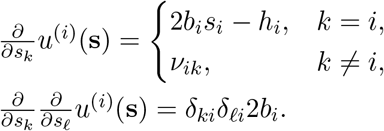

Let **A** be the infinitesimal generator and **M**(*t*) be the mean matrix as defined in Section 2.3. Since

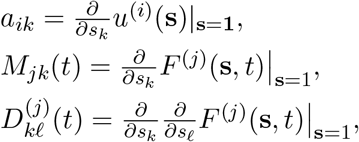

and *u*^(*i*)^(**1**) = 0, we can conclude from (24) that

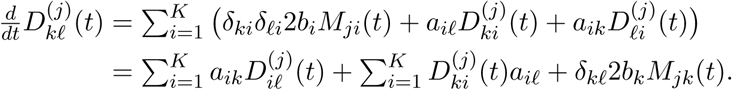

In the second step, we use that 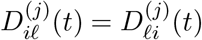. This yields a Lyapunov matrix differential equation,

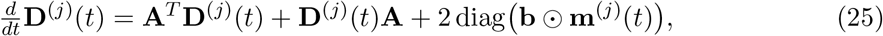

with initial condition **D**^(*j*)^(0) = **0**. The solution is

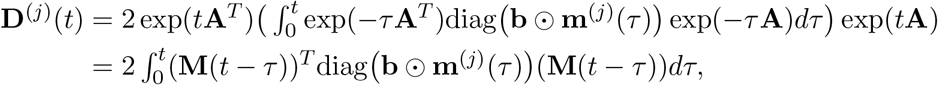

and the expression (4) for **Σ**^(*j*)^(*t*) follows from the fact that

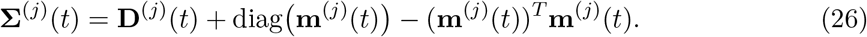

## F Proof of Proposition 2

*Proof of Proposition 2*. Recall from (22) in the proof of Proposition 1 that we can write

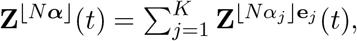

where 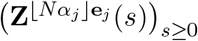 for *j* = 1, …, *K* are independent branching processes started with *⌊Nα*_*j*_ *⌋* cells of type-*j*, respectively. Define

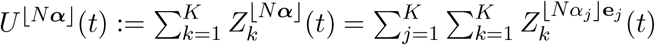

as the total population size at time *t* and note that

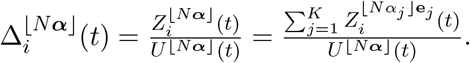

We can therefore write

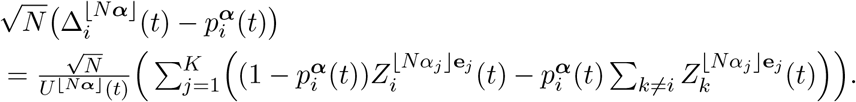

Note that by definition,

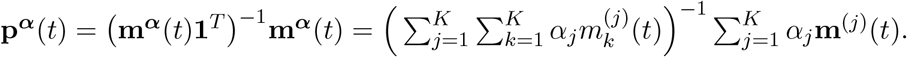

It follows that

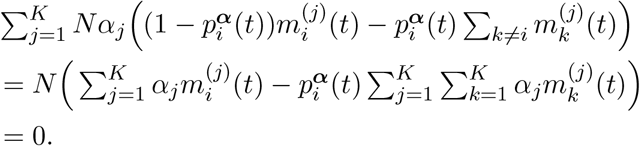

We can therefore write

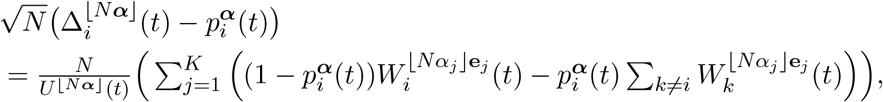

where the vector 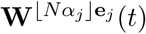 is defined as in (23). In vector form, this becomes

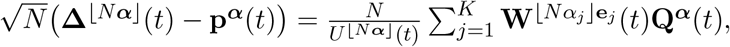

where **Q**^***α***^(*t*) is defined as in (10). By the law of large numbers, *U* ^*N****α***^ (*t*)*/N* → **m**^***α***^(*t*)**1**^*T*^ almost surely as *N* → ∞. Let *J* ≥ 1 and let **C** be a *K × J* matrix. By the (multivariate) central limit theorem,

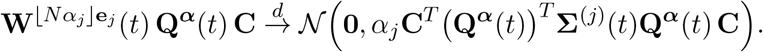

Writing 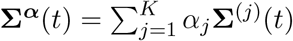, it finally follows from Slutsky’s theorem that

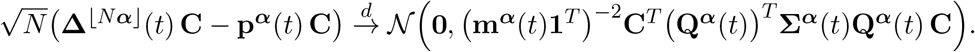

It remains to show that **p**^***α***^(*t*) can be written solely as a function of the switching rates ***ν*** and the net birth rate differences ***λ***^[−*j*]^ for any *j* = 1, …, *K*. To this end, we define

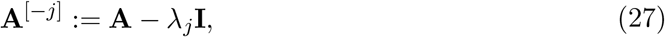

where **I** is the *K × K* identity matrix, and

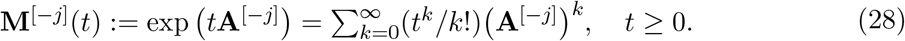

Note that **A**^[−*j*]^ and **M**^[−*j*]^(*t*) only depend on ***ν*** and ***λ***^[−*j*]^. It is easy to see that

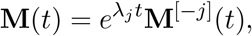

from which it follows that

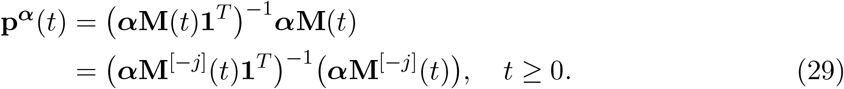

This completes the proof.

## G Proof of Proposition 3

*Proof of Proposition 3*.

1. Since 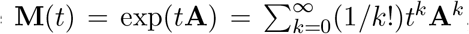, we have 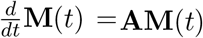. By taking *t* = 0 and noting that **M**(0) = **I**, we obtain

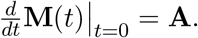 If 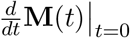 is known, we can recover the switching rate *ν*_*jk*_ for *k*≠*j* by recalling that *a*_*jk* =_ *ν*_*jk*_. We can then recover *λ*_*j*_ for *j* = 1, …, *K* by recalling that *a*_*jj*_ = λ_*j*_ − Σ _*k*≠*j*_ *ν*_*jk*_.
2. Recall that **m**^(*j*)^(*t*) = **e**_*j*_**M**(*t*). By (26) in the proof of Proposition 1, we can write

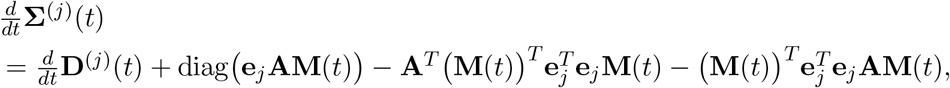

where **D**^(*j*)^(*t*) is the matrix of second factorial moments of **Z**^(*j*)^(*t*). Next, by taking *t* = 0 in (25) and noting that **D**^(*j*)^(0) = **0** and **m**^(*j*)^(0) = **e**_*j*_ for all *j* = 1, …, *K*, we see that

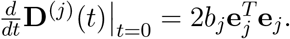

It follows that

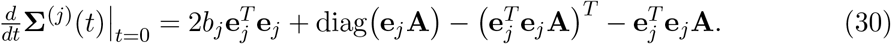

For each *j* = 1, …, *K*, if the switching rates *νjk* for *k* ≠ *j* and the net birth rate *λ*_*j*_ are known, the birth rate *b*_*j*_ can be recovered from 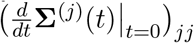 using this expression.

## H Proof of Proposition 4

*Proof of Proposition 4*. We begin by establishing some notation. First, define 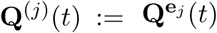 and **Q**^(*j*)^ := **Q**^(*j*)^(0) = **I** − **1**^*T*^ **e**_*j*_, with 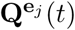 defined as in (10). Also define

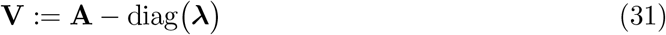

as the infinitesimal generator **A** with the net birth rates ***λ*** removed from the diagonal. Let **v**^(*j*)^ and note that denote the *j*-th row vector of **V** with coordinates 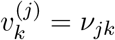 for *k* ≠ *j* and 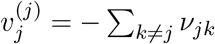 and note that

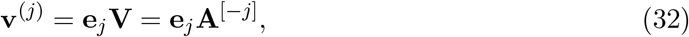

where **A**^[−*j*]^ is defined as in (27). Also note that **v**^(*j*)^**1**^*T*^ = 0. In the proof, we will rely on the following basic facts:

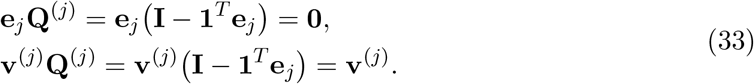

1. Since **p**^(*j*)^(*t*) = **e**_*j*_ exp(*t***A**)**1**^*T*^ −1 **e**_*j*_ exp(*t***A**), we can write

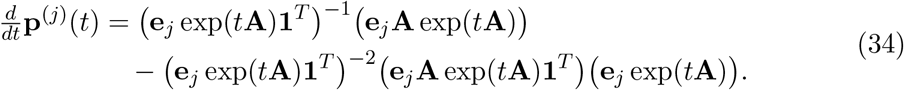

Since exp(**0**) = **I, e**_*j*_**1**^*T*^ = 1 and **e**_*j*_**A1**^*T*^ = *λ*_*j*_, we obtain by (32),

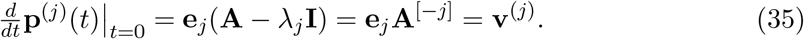

Since the *k*-th coordinIate of **v**^(*j*)^ is *ν*_*jk*_ for *k* ≠ *j*, we can recover *ν*_*jk*_ from the *k*-th coordinate of 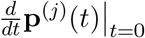.
2. i. Using (34), we begin by writing

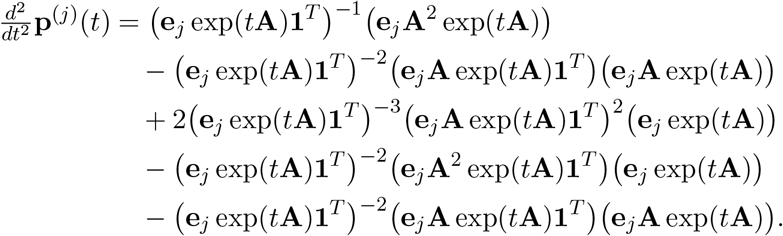

Since exp(**0**) = **I, e**_*j*_**1**^*T*^ = 1, **e**_*j*_**A1**^*T*^ = *λ*_*j*_, **v**^(*j*)^ = **e**_*j*_**A**^[−*j*]^ and **Q**^(*j*)^ = **I** − **1**^*T*^ **e**_*j*_,

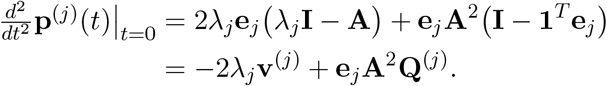

Recalling that **A** = **A**^[−*j*]^ + *λ*_*j*_**I** by (27), we can write

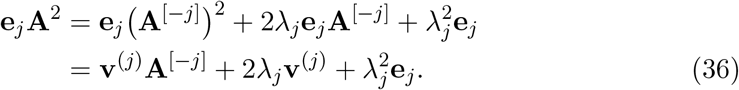

Since **e**_*j*_**Q**^(*j*)^ = **0** and **v**^(*j*)^**Q**^(*j*)^ = **v**^(*j*)^ by (33), it follows that

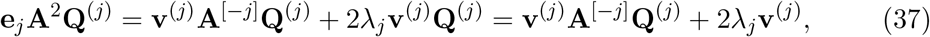

which implies

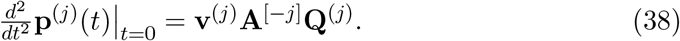

It is straightforward to verify that for *i* ≠ *j*,

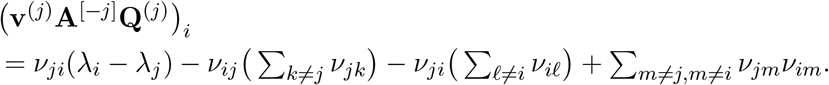

If ***ν*** and 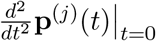 are known, we can therefore use (38) to get an equation for *λ*_*i*_ − *λ*_*j*_ of the form *ν*_*ji*_(*λ*_*i*_ − *λ*_*j*_) = *C* for some constant *C*. If *ν*_*ji*_ 0, we immediately obtain the value of *λ*_*i*_ − *λ*_*j*_. If *ν*_*ji*_ = 0, then by our assumption of irreducibility, there exist integers *n*_1_, …, *n*_*k*_ so that 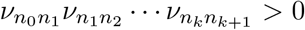, where *n*_0_ = *j* and *n*_*k*+1_ = *i*. For each *ℓ* = 0, …, *k*, we can use the fact that 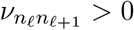 to obtain the value of 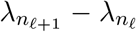. Since 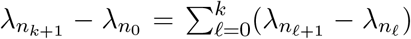,we also obtain the value of *λ*_*i*_ − *λ*_*j*_.
ii. We know from (34) that

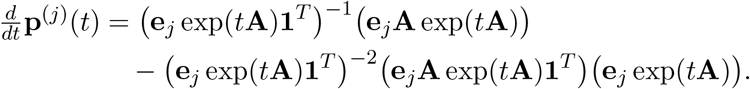

We also know from (1) that

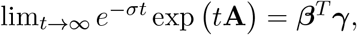

where ***β*** and ***γ*** are positive vectors. It follows that as *t* → ∞,

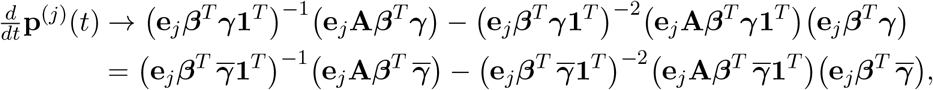

where 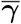 is the normalized version of ***γ***, see (2). Since **e**_*j*_***β***^*T*^ = *β*_*j*_ *>* 0 and 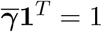, we obtain

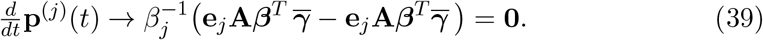

On the other hand, by noting that **A** and exp(*t***A**) commute, we can rewrite the expression (34) for 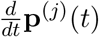 as

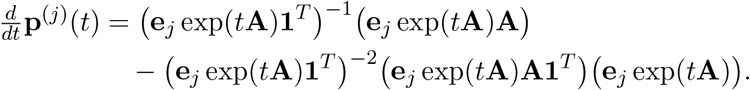

Since **A1**^*T*^ = ***λ***^*T*^, **A** = **V**+diag(***λ***) by (31), 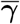 diag(***λ***) = ***λ*** diag 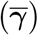 and 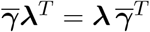, we get as *t* → ∞,

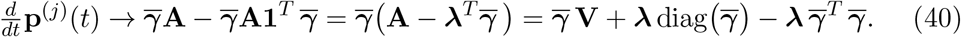

Combining (39) and (40), we obtain the following linear system for ***λ***:

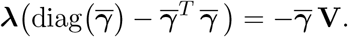

It is straightforward to verify that this system is solved by

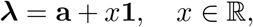

for some vector **a**, which can be used to extract ***λ***^[−1]^.
3. By the definition of **S**^(*j*)^(*t*) in (10),

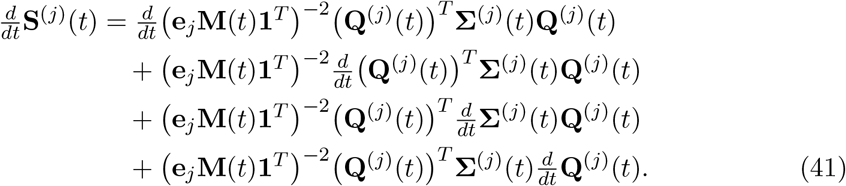

Since **Σ**^(*j*)^(0) = **0** and **e**_*j*_**M**(0)**1**^*T*^ = 1, we obtain

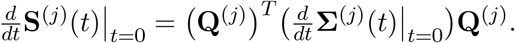

From (30) in the proof of Proposition 3, we know that

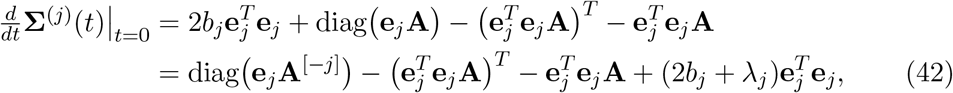

where in the second step, we write **A** = **A**^[−*j*]^ +*λ*_*j*_**I**. Since **e**_*j*_**Q**^(*j*)^ = **0** and **e**_*j*_**A**^[−*j*]^ = **v**^(*j*)^, we obtain

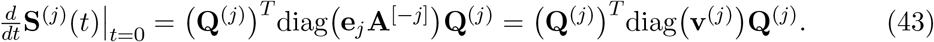

It is straightforward to verify that the (*j, k*)-th coordinate of (**Q**^(*j*))^ *T* diag (**v**^(*j*))^ **Q**^(*j*)^ is −*ν*_*jk*_. Thus, knowledge of the switching rates ***ν*** follows immediately from knowledge of 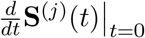 for *j* = 1, …, *K*, but no other parameters can be extracted.

## Acknowledgments

EBG and KL were supported in part by NSF grant CMMI-1552764. JF was supported in part by NSF grants DMS-1349724 and DMS-2052465. KL and JF were supported in part by the Research Council of Norway R&D Grant 309273. EBG was supported in part by the Norwegian Centennial Chair grant and the Doctoral Dissertation Fellowship from the University of Minnesota.

